# McrD binds asymmetrically to methyl-coenzyme M reductase improving active site accessibility during assembly

**DOI:** 10.1101/2023.02.01.526716

**Authors:** Grayson L. Chadwick, Aaron M.N. Joiner, Sangeetha Ramesh, Douglas A. Mitchell, Dipti D. Nayak

## Abstract

Methyl-coenzyme M reductase (MCR) catalyzes the formation of methane and its activity accounts for nearly all biologically produced methane released into the atmosphere. The assembly of MCR is an intricate process involving the installation of a complex set of post-translational modifications and the unique Ni porphyrin cofactor F_430_. Despite decades of research, details of MCR assembly remain largely unresolved. Here, we report the structural characterization of MCR in two intermediate states of assembly. These intermediate states lack one or both F_430_ cofactors and form complexes with the previously uncharacterized McrD protein. McrD is found to bind asymmetrically to MCR, displacing large regions of the alpha subunit and increasing active site accessibility for the installation of F_430_—shedding light on the assembly of MCR and the role of McrD therein. This work offers crucial information for the expression of MCR in a heterologous host and provides new targets for the design of MCR inhibitors.

**One-sentence summary:** Structural characterization of methyl-coenzyme M reductase assembly intermediates.

## Main text

Methyl-coenzyme M reductase (MCR) is the enzyme responsible for nearly all biological methane production and is found exclusively in the domain Archaea. MCR is a C2-symmetric heterohexamer defined by α_2_β_2_γ_2_ subunit stoichiometry (**fig. S1A**). Substrate tunnels lined with post-translational modifications (PTMs) end in two active sites formed at the interface of the α, α’, β and γ subunits and α’, α, β’ and γ’ on the opposing side (**fig. S1B-C**). Each active site contains F_430_, a unique Ni porphinoid that catalyzes the reduction of methyl-coenzyme M (CH3-CoM) by coenzyme B (CoB), yielding methane and a heterodisulfide (*1*) (**fig. S1D**). Subsequent studies have determined MCR structures from diverse methanogens (*2*), mutants lacking specific substrate channel PTMs (*3, 4*), anaerobic methanotrophic (ANME) archaea, which use the enzyme in the reverse direction for methane oxidation (*5*), and the recently discovered Alkyl-coenzyme M reductases (ACR), which can oxidize short chain alkanes (*6*). Together, this research has facilitated studies on the reaction mechanism of MCR (*7*), and the mode of action for MCR inhibitors used to reduce the emission of this potent greenhouse gas (*8, 9*).

Despite the advances in our understanding of MCR structure and function, key questions regarding its assembly and activation remain unresolved. For instance, the three subunits αβγ encoded by the *mcrABG* genes are found together in an operon with two additional genes: *mcrC* and *mcrD* (**Fig. 1A**). The function of the McrC and McrD proteins is unknown but they are universally conserved in methanogens and have been suggested to be involved in MCR activation (*10*) and assembly (*11*), respectively. McrD associates with MCR in native organisms (*12*) and when heterologously expressed (*11*), and McrD produced in *E. coli* stimulated the final step of F_430_ biosynthesis(*13*). Based on these observations, McrD is hypothesized to be a chaperone required for the insertion of F_430_ into a previously unobserved apo-MCR (*11, 14*). Here, we utilize Cas9-based genome editing in *Methanosarcina acetivorans* (*15*) to characterize McrD and employ single-particle cryoelectron microscopy (cryoEM) to visualize McrD-bound MCR assembly intermediates.

**Fig. 1.**
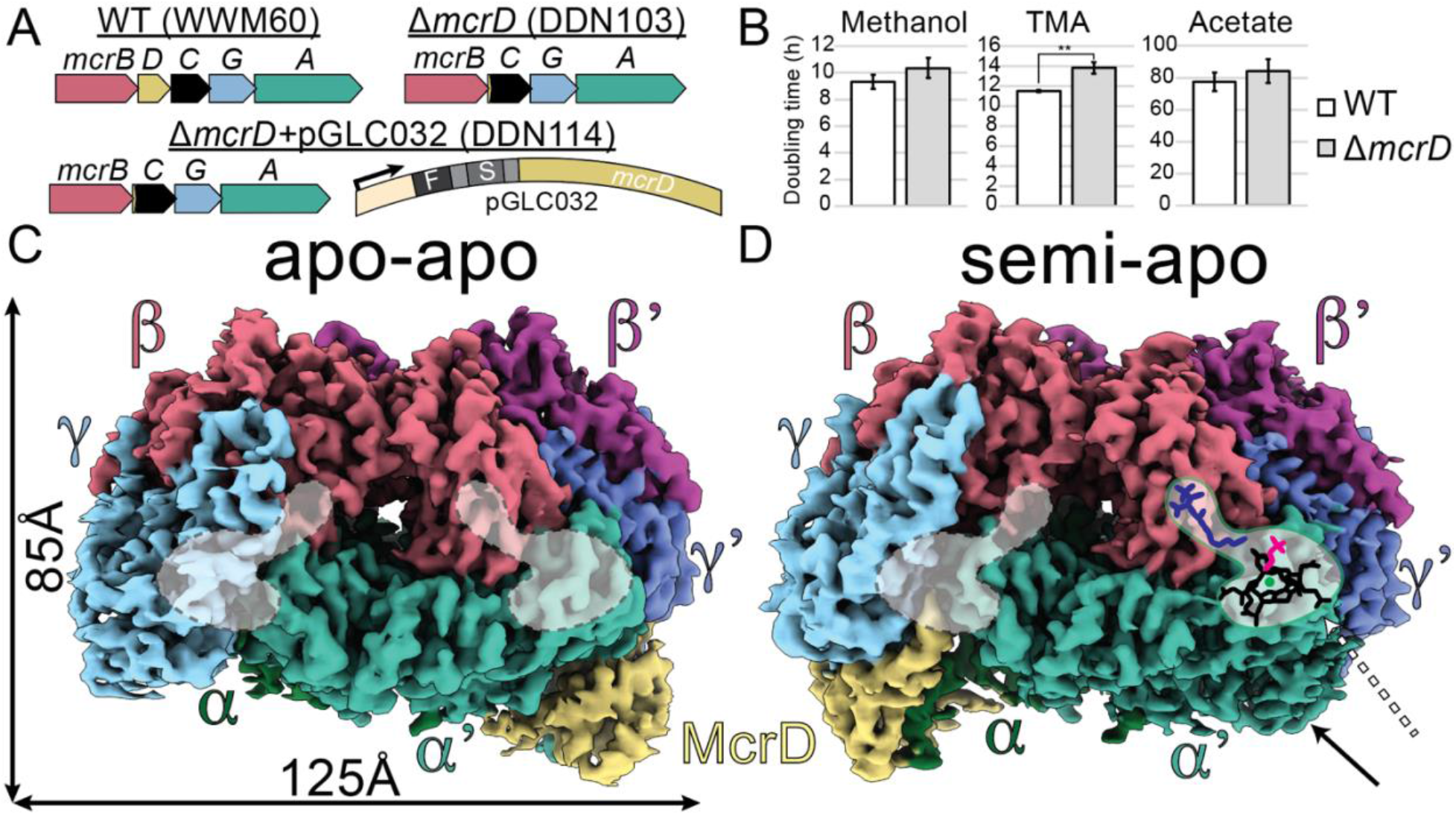
Construction of *ΔmcrD* strain and cryoEM structure of McrD-bound MCR complexes. (**A**) MCR operon in the parental strain (WWM60), *ΔmcrD* mutant (DDN103), and tagged complementation strain (DDN114). Arrow indicates pMcrB(tetO1) promoter used for tetracycline inducible expression of McrD, F and S represent FLAG and Strep tags, respectively. (**B**) Doubling times of WT and *ΔmcrD* strain on methanol (MeOH), trimethylamine (TMA) and acetate (** p-value < 0.01, two-sided t-test). (**C, D**) CryoEM densities of the apo-apo and semi-apo McrABGD complexes (**Supplementary Table 4)**. Empty active sites are outlined with dotted lines, while filled active site on the contralateral side of the semi-apo model is indicated by the dashed arrow. Solid arrow indicates density corresponding to the N-terminal domain of the α’ subunit in the semi-apo complex that is absent in the apo-apo complex.

Surprisingly, although the entire MCR operon is essential (*16*), the *mcrD* gene is not in *M. acetivorans* (**Fig. 1A**). We generated a *ΔmcrD* mutant and verified the absence of compensatory mutations or off-target editing by whole genome resequencing (**fig. S2**). This *ΔmcrD* strain exhibited only a minor increase in doubling time compared to the parental strain when grown on a variety of substrates (**Fig. 1B**). The growth defect remained negligible in cells stressed by nickel limitation, oxygen exposure and non-ideal growth temperatures (**fig. S3**). Consistent with this minor growth defect, only a small number of genes were significantly differentially expressed in the *ΔmcrD* mutant (**fig. S4**).

We used a tetracycline-inducible expression vector to reintroduce the *mcrD* gene with N-terminal tandem FLAG and Strep tags into the *ΔmcrD* background (**Fig. 1A**). McrD expression was verified by Western blot against the FLAG epitope in crude cell extracts (**fig. S5**). Affinity purification of tagged McrD yielded stoichiometric amounts of McrA, B and G subunits, indicative of a McrABGD complex in *M. acetivorans* (**fig. S6**). This complex is less thermostable and exhibited far less F_430_ absorbance compared to full assembled MCR (*17*) (**fig. S7**). Intriguingly, this complex could only be recovered from actively dividing cells, while purification from cells in stationary phase yielded predominantly free McrD (**fig. S6**). The presence of an McrABGD complex primarily in exponential phase is consistent with its role in the assembly of MCR rather than in a repair pathway to salvage damaged MCR. Additionally, isolation of the McrABGD complex was insensitive to the location of affinity tags on McrD, suggesting that it forms through a biologically relevant interaction between McrD and the other subunits, not a spurious interaction between the tag and MCR (**fig. S6**).

Two distinct conformational states of the McrABGD complex were discovered by cryoEM: 1) semi-apo at 3Å and 2) apo-apo at 3.1Å (**Fig. 1C, D**, **fig. S8–9**). We docked a previous crystal structure, of the MCR_ox1-silent_ state purified from *Methanosarcina barkeri* (*2*), along with an AlphaFold2 (*18,19*) prediction of McrD into both densities and then manually rebuilt each model. Surprisingly, all of the cryoEM density in both maps could be accounted for by a complex with α_2_β_2_γ_2_D1 stoichiometry. These asymmetric complexes were supported by a molar mass estimate, acquired by size exclusion chromatography coupled with multi-angle light scattering, of roughly 256kDa, which is too small for α_2_β_2_γ_2_D2 (~313kDa) and too large for α_1_β_1_γ_1_D_1_ (~157kDa) (**fig. S10**). Consistent with the hypothesis that McrD assists in the formation of an assembly intermediate, the two conformations differ in their active site occupancy. Both active sites in the apo-apo model are empty whereas the semi-apo model contains one empty active site and the other one shows clear density corresponding to the presence of three ligands (F_430_, CoM, and CoB) (**Fig. 1C, D**).

In each conformational state, McrD is nestled between the γ, α and α’ subunits (semi-apo lettering, **Fig. 2A**). Residues 10-129 of McrD are resolvable in our cryoEM density and adopt aβαββαβ fold motif. A large loop between the two βαβ repeats of McrD, composed of residues 47-68, extends upwards forming a wedge between the γ and α’ subunits, positioning the bulk of McrD in a region normally occupied by the N-terminal domain of the α subunit (**Fig. 2B**). The N-terminal domain of the α subunit bends down and outwards along the β-sheet of McrD, converting Leu^α78^-Arg^α81^ into a β-strand (**Fig. 2B**). Although the C-terminus of McrD is partially unresolved in our structures, overlaying the full-length AlphaFold2 prediction reveals that the C-terminal domain contributes to the formation of a small putative tunnel into which the Leu^α78^-Arg^α81^β-strand fits neatly (**fig. S11**). The formation of this β-strand involves a major reorganization of the N-terminal domain of the α subunit (**Fig. 2C**). In both conformations, binding of McrD results in large-scale movements of γ and β compared to previous crystal structures (**Fig 2D**), resulting in increased accessibility of the active site on the McrD-bound side. The resolution of the cryoEM structures was sufficient to visualize the methylations of Arg^285^ and Cys^472^ in α and α’ (**Fig. 2E**), which were also detected by high-resolution and tandem mass spectrometry (HRMS/MS) (**fig. S12–S14**). The didehydro Asp^470^, thio Gly^465^ and *N-*methyl His^271^ modifications were detected and verified by HRMS/MS despite ambiguity in the cryoEM data (**Fig. 2F**).

**Fig. 2.**
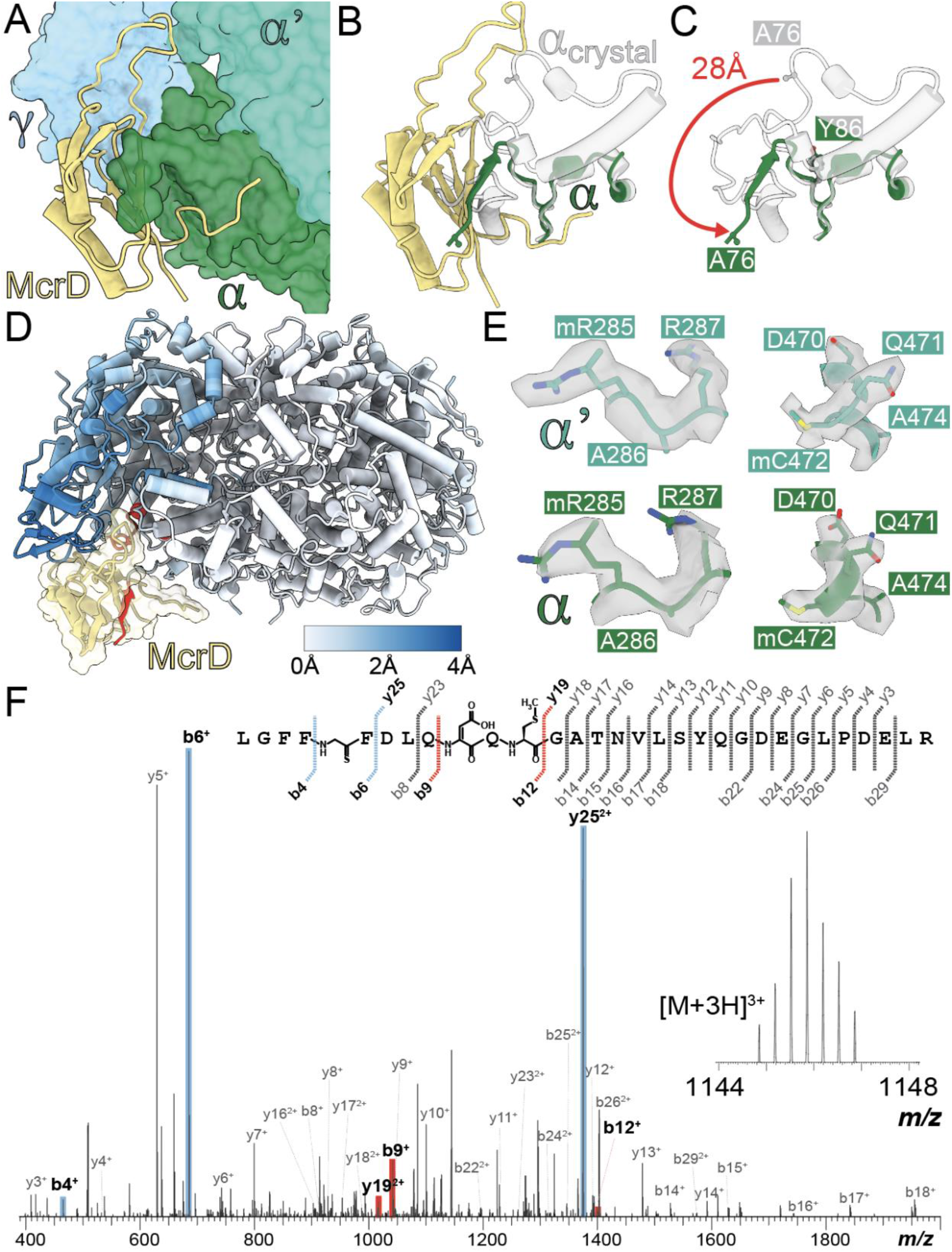
McrD-binding to PTM-bearing MCR causes large-scale structural changes. (**A**) Detail of McrD binding location on the semi-apo model. (**B**) The α subunit N-terminal domain from the *M. barkeri* MCR crystal structure (PDB: 1e6y) depicted in white, with the corresponding region of the semi-apo α subunit dark green. (**C**) Significant rearrangement exemplified by Ala^α76^ which deviates 28Å upon McrD binding. All residues before Ala^α76^ are disordered. (**D**) Ribbon diagram of semi-apo structure colored by Cα deviations from 1e6y. Residues deviating by more than the color scale maximum of 4Å are shown in red. (**E**) Density corresponding to methylations at Arg^285^ and Cys^472^ in the α and α’ subunits. (**F**) HRMS/MS fragmentation and annotated collision-induced dissociation spectrum of an McrA tryptic peptide (Leu^461^-Arg^491^) conclusively identified three of the five PTMs. Cyan and red mass peaks indicate the most diagnostic ions for localizing the modification sites. For additional details and HRMS/MS data confirming the remaining PTMs, see **fig. S12–S14**.

The asymmetric nature of these structures results in four unique active site conformations (**Fig. 3A,B**). In both models, the active site on the McrD-bound side is the most distorted compared to the crystal structure of fully assembled MCR (**Fig. 3C, S15A**). Nearly all active site forming secondary structures deviate significantly and are unresolvable in some places, most notably the loops between Leu^α333^-Tyr^α346^ and Ala^α158^-Glu^α166^ containing the F_430_ axial ligand Gln^α161^. The loop between Asp^α414^ and Phe^α416^ show a dramatic rearrangement, with His ^α415^ and Phe ^α416^ moving in to occupy regions normally filled by F_430_ and the axial ligand loop. Additional structural changes include a distortion to the loop containing Tyr^β365^ resulting in a reorientation of the hydroxy group by 180 degrees.

**Fig. 3.**
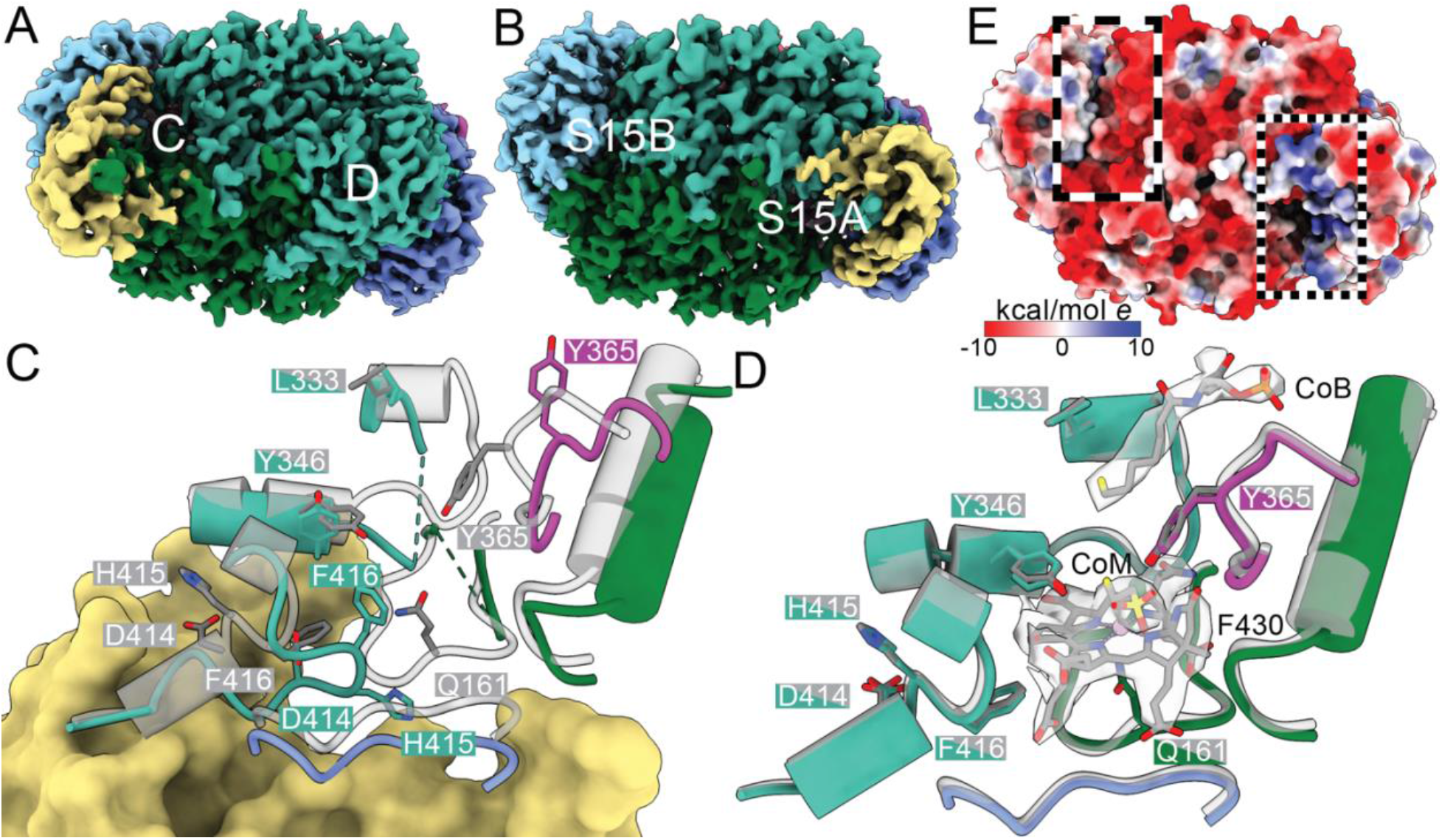
Active site variations in MCR assembly intermediates. (**A-B**) Bottom view of complexes with labels showing active sites detailed in subsequent panels. (**C-D**) Active site details of the semi-apo model with key residue side chains shown (for corresponding figures for apo-apo model see **fig. S15**). MCR crystal structure from *M. barkeri* (PDB:1e6y) overlaid in gray. In D cryoEM density corresponding to CoM, CoB and F_430_ is shown in transparent gray. (**E**) Bottom view of the apo-apo complex colored by electrostatic potential. Boxed regions on the McrD-bound side (dots) and contralateral side (dashes) highlight strong electrostatic differences, with positively charged patches leading to the active site from McrD and the α’ subunit. Boxes indicate similar positions on the side opposite McrD with the axial ligand loop blocking the active site and mostly negatively charged surfaces.

The contralateral active sites differ significantly between the semi-apo and apo-apo states. In the semi-apo complex, the contralateral active is nearly identical to those observed in the fully assembled crystal structure (**Fig. 3D**). Strong density is present throughout the entire active site cavity, allowing for unambiguous identification of CoM, CoB and F_430_ as well as the loop containing Gln^α161^. Here, the N-terminal region of the α’ subunit caps the active site as in the crystal structure. In stark contrast, the contralateral active site of the apo-apo structure displays weak density for the loop with Gln^α161^, and lacks distinct density for the ligands and the entire N-terminal domain of the α’ subunit (**fig. S15B**). There is also a prominent deformation of the helix-loop-helix region between Leu^α’333^ and Tyr^α’346^ that could not be fully resolved. Notably, Gln^α161^ has shifted ~7Å away from the active site cavity, consistent with a lack of F_430_. A comparison of the two active sites in the apo-apo model highlights the importance of McrD binding for F_430_ ingress to the active site. The disordered axial ligand loop leads to a solvent-exposed pathway into the active site that is lined with basic residues contributed by both McrD and the β-strand Leu^α’78^-Arg^α’81^ (**Fig. 3F**). In contrast, the contralateral active site remains far less accessible due to a surface covered mostly by acidic residues and an axial ligand loop left largely in place.

Based on these data, we propose a model for the assembly of the MCR complex and the role of McrD in this process (**Fig. 4**). Our model begins with a PTM-containing apo-α_2_β_2_γ_2_ MCR that is folded except for the α subunit N-terminal domains. McrD is not needed to destabilize the α N-terminal domains, as evidenced by the unstructured contralateral N-terminus in our apo-apo α_2_β_2_γ_2_D_1_ structure. Free McrD recognizes one of the unfolded N-terminal domains to bind to the complex, yielding our observed apo-apo α_2_β_2_γ_2_D_1_ state. Though they have never been visualized, there are two hypothetical ways that assembly might proceed: 1) the association of a second McrD on the contralateral side leading to an apo-apo α_2_β_2_γ_2_D_2_ complex, or 2) the insertion of F_430_, dissociation of McrD, and folding of the alpha N-terminal domain to cap the active site producing a semi-apo α_2_β_2_γ_2_ state. While our purification strategy would never capture the latter state, it should have trapped the former, if present in significant quantities. In the first case, F_430_ insertion followed by McrD loss, or in the second case, rebinding of McrD to the opposite side, would lead to our observed semi-apo α_2_β_2_γ_2_D_1_. From here, an additional round of F_430_ insertion, loss of McrD, and folding of the alpha N-terminal domain to cap the active site would yield a conformation identical to the crystallized MCR_ox1-silent_ form, which is ready to be a substrate for the activation complex.

**Fig. 4.**
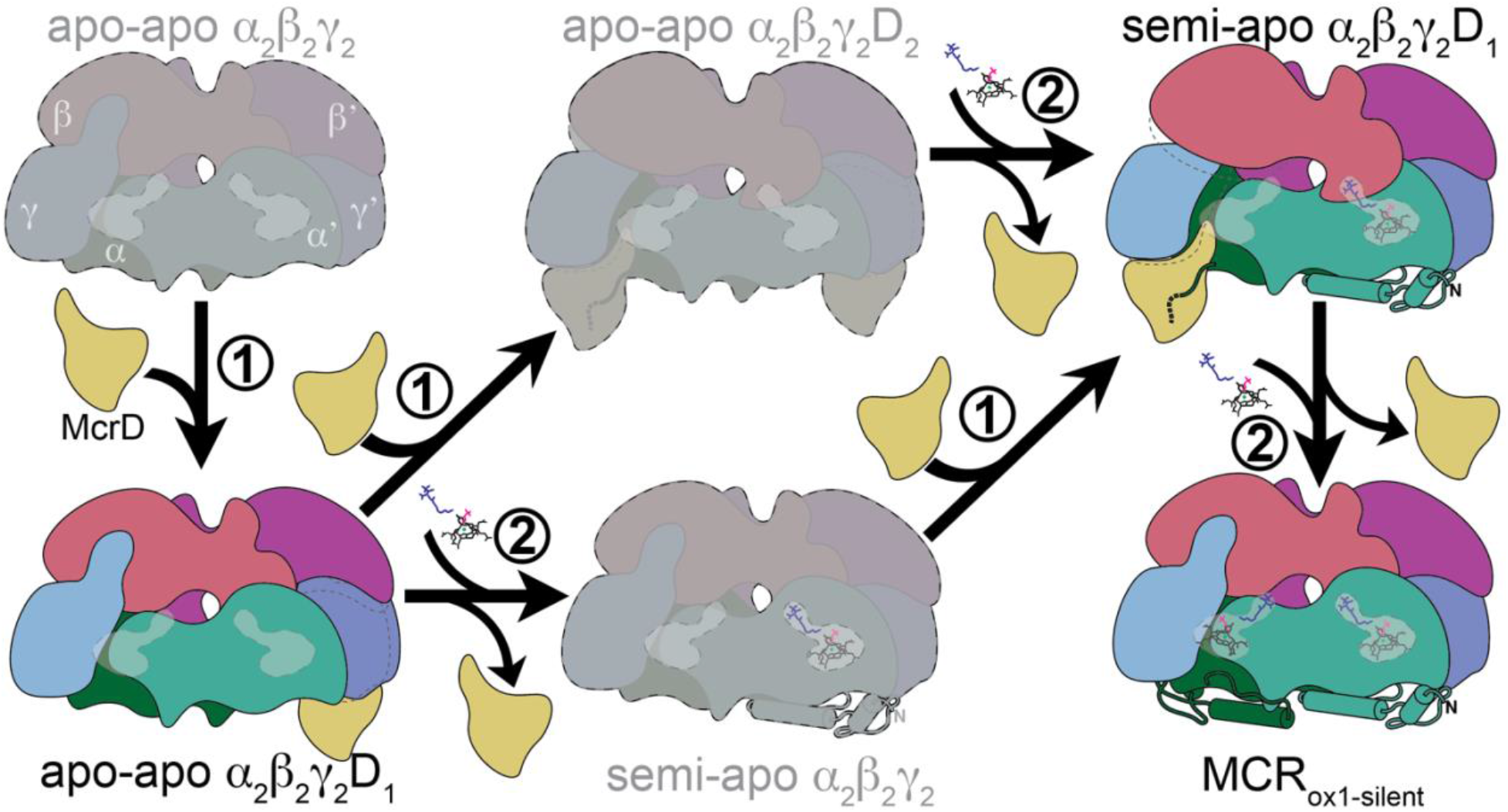
Role of McrD in MCR assembly. Experimentally observed complexes are shown in full color, while hypothesized intermediates are shown in gray. The two proposed functions of McrD are numbered. 1) McrD recognizes the disordered N-terminal domain of an α subunit lacking F_430_, and 2) McrD and possibly unidentified additional factors facilitate the insertion of F_430_, McrD dissociation and α subunit N-terminal domain folding. The order of these two functions is not clear, leading to two possible routes between our observed assembly intermediates.

Here we have provided the first structural insight into the assembly of MCR and McrD’s role therein. Similar to the PTMs on the alpha subunit (*3*), the loss of McrD has little if any growth penalty under a variety of laboratory conditions, highlighting that important facets of MCR biology in nature not captured by ideal laboratory conditions. Despite this, our structural insight into empty MCR active sites could propel the design of novel anti-methanogen compounds to block F_430_ insertion. Potent compounds that inhibit MCR, particularly as additives to ruminant feedstocks (*20*), will likely be a reliable and cost-effective strategy to curb methane emissions, which is a critical aspect of international efforts to limit global temperature rise. A complete understanding of the MCR enzyme’s assembly, activation and reaction mechanism is crucial for the development and deployment of diverse classes of MCR antagonists. Furthermore, elucidating the process of MCR assembly and activation advances the biotechnological goal of heterologous expression of the methane generating metabolic pathway, and contributes to our comprehension of the evolutionary transitions required for the natural horizontal transfer of this metabolism (*21*). Future work must identify what additional cellular factors, if any, associate with the McrD-bound assembly intermediates to facilitate F_430_ insertion and N-terminal domain folding.

## Materials and Methods

See supplemental information.

## Acknowledgements

Funding list for DDN: Beckman Young Investigator Award sponsored by the Arnold and Mabel Beckman Foundation, Searle Scholars Program sponsored by the Kinship Foundation, the Rose Hills Innovator Grant, the Packard Fellowship in Science and Engineering sponsored by the David and Lucille Packard Foundation, and the Simons Early Career Investigator in Marine Microbial Ecology and Evolution Grant sponsored by the Simons Foundation. DDN is a Chan Zuckerberg Biohub Investigator. GLC and AMNJ are supported by the Miller Institute for Basic Research in Science, University of California Berkeley. This work was supported in part by a grant from the National Institutes of Health (GM097142 to D.A.M.).

## Supplementary Information

**fig. S1.**
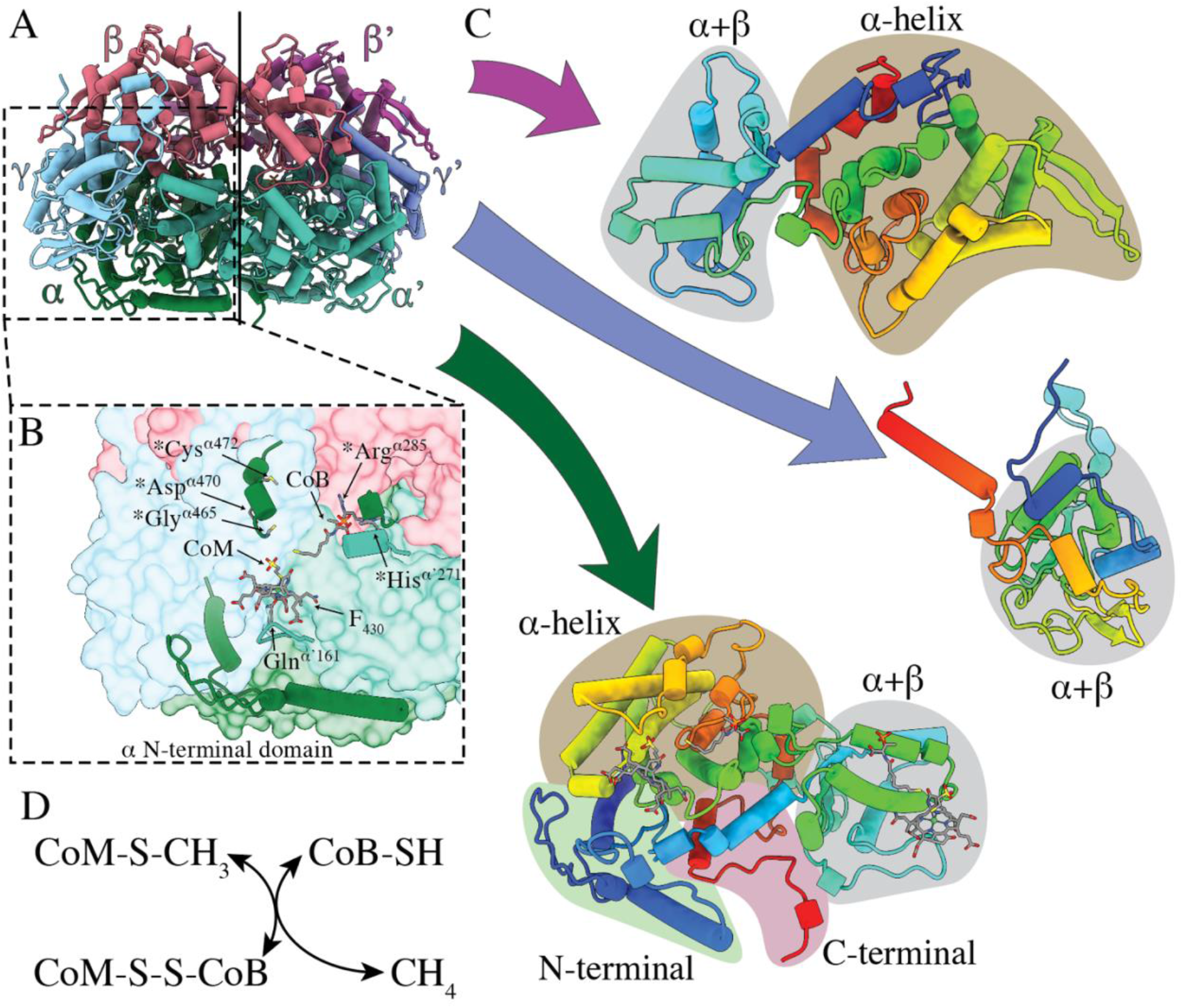
MCR reaction and structural features. (**A**) Crystal structure of methyl-coenzyme M reductase (MCR) from *M. barkeri* demonstrating the quaternary structure of the complex (PDB:1e6y). (**B**) Detail of MCR active site showing coenzyme M (CoM), coenzyme B (CoB), cofactor F_430_, F_430_’s axial ligand Gln^α^’^161^ and the five post-translationally modified residues denoted by asterisks. The N-terminal domain of the α subunit that covers the bottom of active site shown as a cartoon. (**C**) Folds of McrB (top), McrG (right) and McrA (bottom) with major domains labelled as defined by Ermler *et al*. 1997. The α-helix and α+β domains of McrA and McrB are structurally similar, suggesting an ancient duplication event. (**D**) The reversible reaction catalyzed by MCR. Methyl coenzyme M (CoM-S-CH3) is reduced by coenzyme B (CoB-SH) to form the heterodisulfide (CoM-S-S-CoB) and methane.

**fig. S2.**
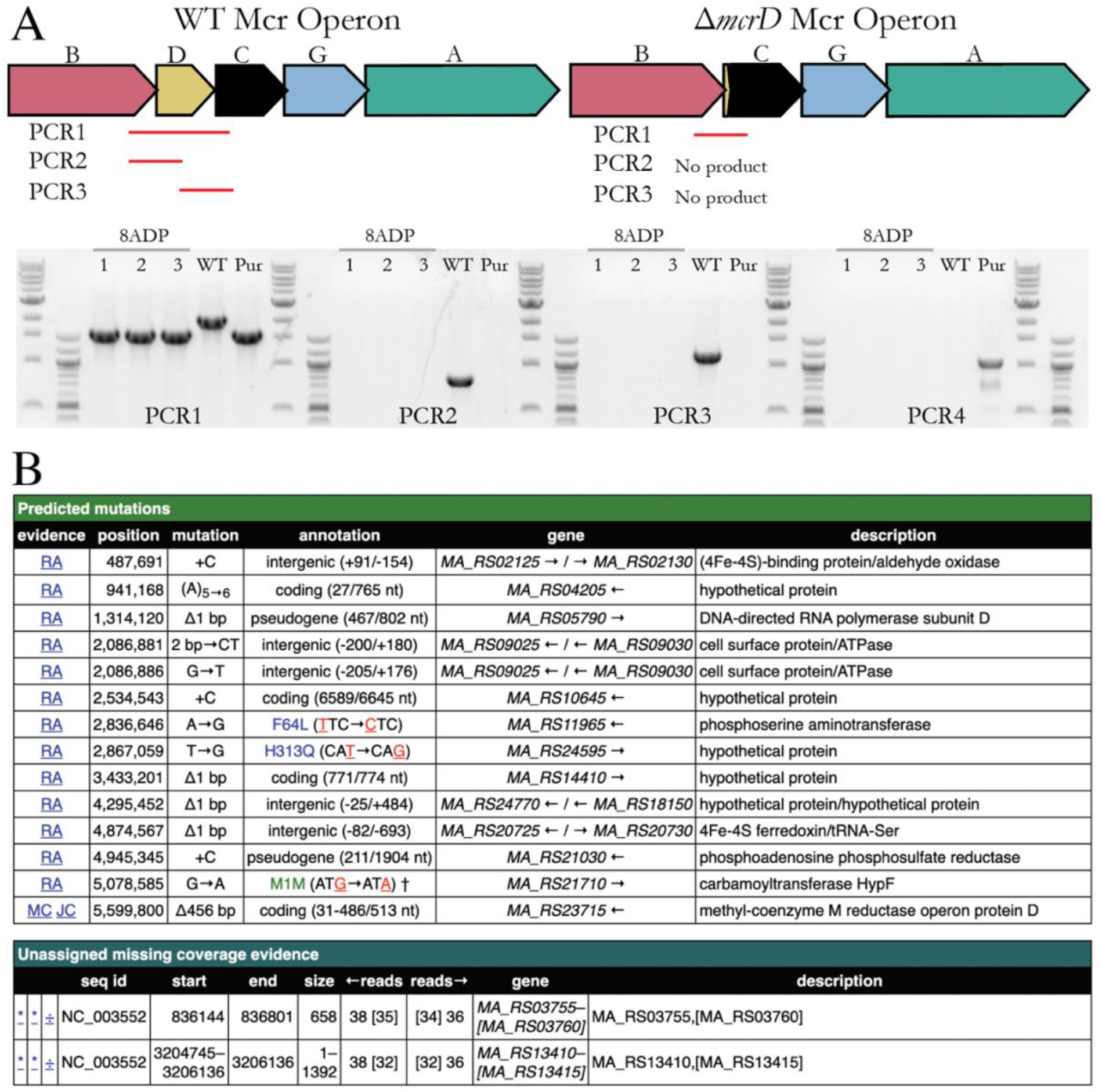
Deletion of the *mcrD* gene. (**A**) PCR reactions spanning the *mcrD* gene, as well as primers within the *mcrD* coding sequence were used to verify *mcrD* deletion and rule out wild type contamination. Genomic DNA was used as a template from wildtype (WT; WWM60), initial colony selected from Puromycin plate that contains the gene editing plasmid (Pur), and three colonies from 8-aza-2,6-diaminopurine (8ADP) containing plates used for counterselection against the gene editing plasmid. PCR1 spans the *mcrD* gene, showing a full-sized product only in WT and expected truncated size post gene editing. Similarly, only WT generates a product in PCR 2 and 3 with primers binding within *mcrD*. PCR4 amplifies the *pac* gene conferring puromycin resistance on the gene editing plasmid, verifying plasmid curing by 8ADP counterselection (Nayak and Metcalf 2017). (**B**) Breseq comparison of DDN103 to *M. acetivorans* C2A reference genome. All mutations and missing coverage are identical to the parental strain WWM60 (Nayak and Metcalf 2017) besides the 456 bp deletion of *mcrD*(MA_RS23715). RA: read alignment, MC: missing coverage, JC: new junction.

**fig. S3.**
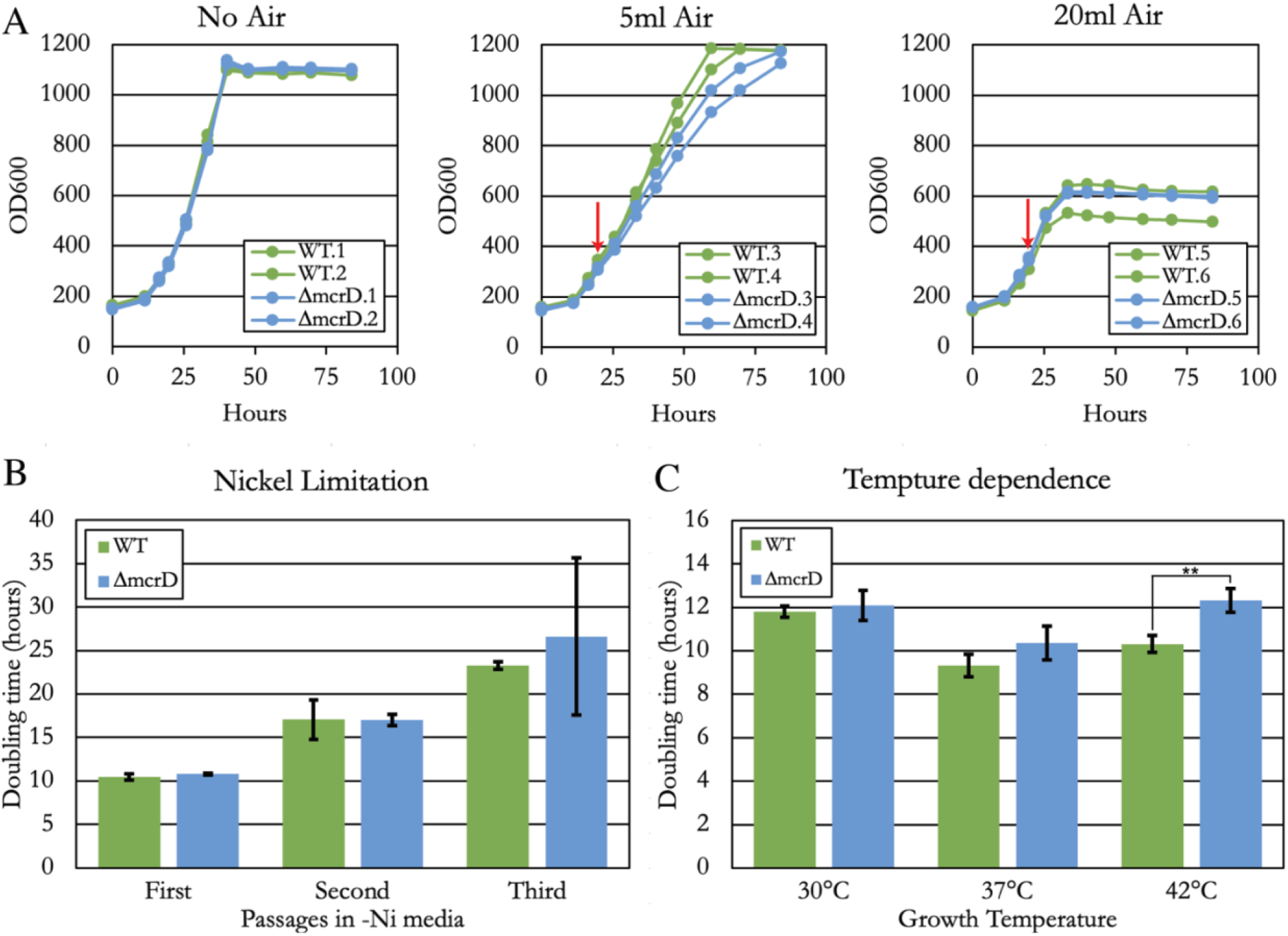
Stress experiments. Attempts to find conditions with a dramatic growth difference between wildtype (WT; WWM60) and *ΔmcrD* were unsuccessful. (**A**) MCR is known to be oxygen sensitive, so air was introduced into growing cultures of WT and *ΔmcrD* at the times indicated by red arrows. Low (5 ml) and high (20 ml) air challenges negatively affected growth, but with no noticeable difference between strains. (**B**) Cultures were washed three times and transferred into methanol media lacking added Ni. Although subsequent passages showed an increase in doubling time for both WT and Δ*mcrD*, no significant difference was observed between strains (two-sided t-test). (**C**) Growth in methanol media in non-ideal low (30°C) and high (42°C) temperatures was assessed. In neither case was the Δ*mcrD* dramatically worse than WT, although at 42°C there was a statistically significant difference (p-value < 0.01, two-sided t-test). Error bars in **B** and **C** represent standard deviations of triplicate cultures.

**fig. S4.**
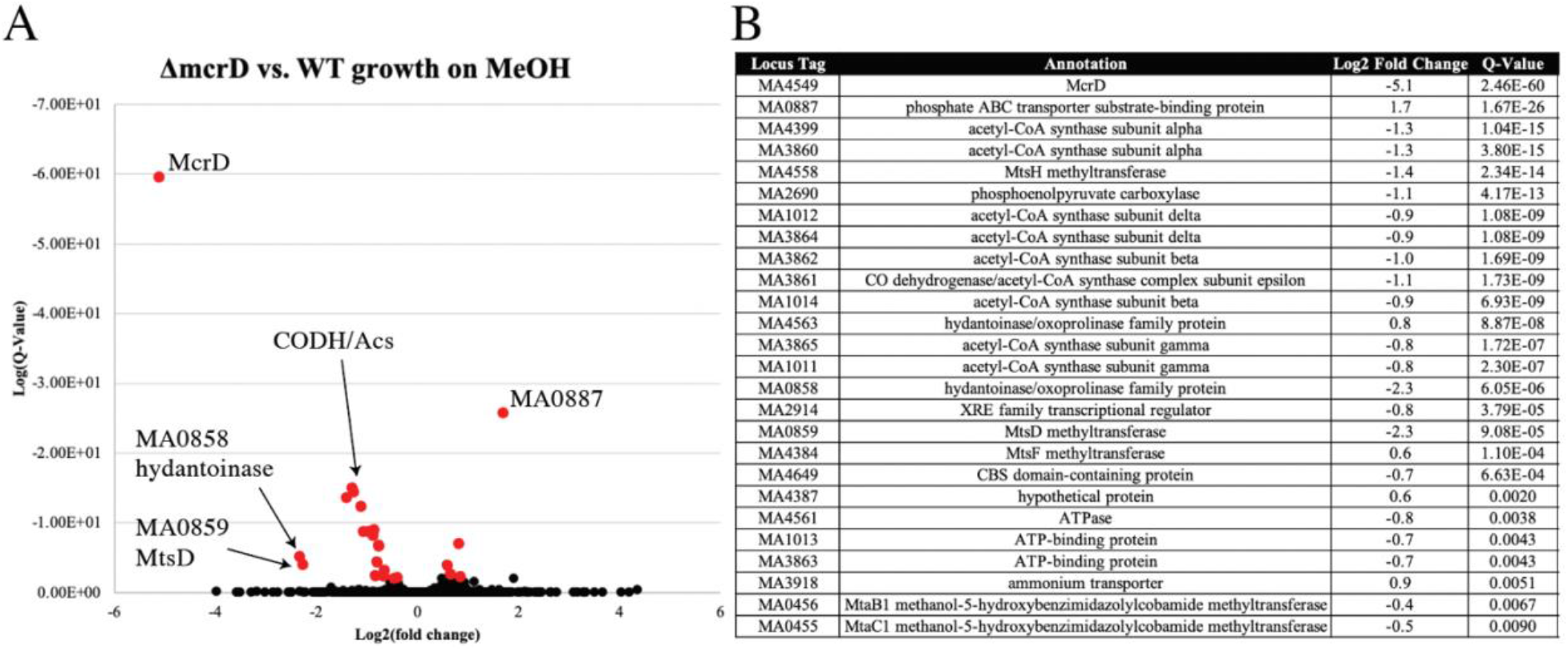
Transcriptome comparison of *ΔmcrD* and WT. (**A**) Comparison of significantly differentially expressed genes in *ΔmcrD* and WT grown on methanol. Only 25 genes aside from *mcrD* were differentially expressed between the two strains (defined as a Q-value<0.01 highlighted in red). There is a downregulation of some carbon metabolism genes including multiple isoforms of CODH/Acs and PEP carboxylase, perhaps suggestive of a decrease in carbon flow towards biomass production. All fold changes are relatively minor, at or below two-fold difference. Importantly, other MCR genes are not differentially expressed, revealing that the absence of the *mcrD* gene has not affected the stability of the MCR operon transcript, or resulted in a transcriptional change in this operon. (**B**) Locus tags, annotations log2 fold change and Q-values for all 25 genes.

**fig. S5.**
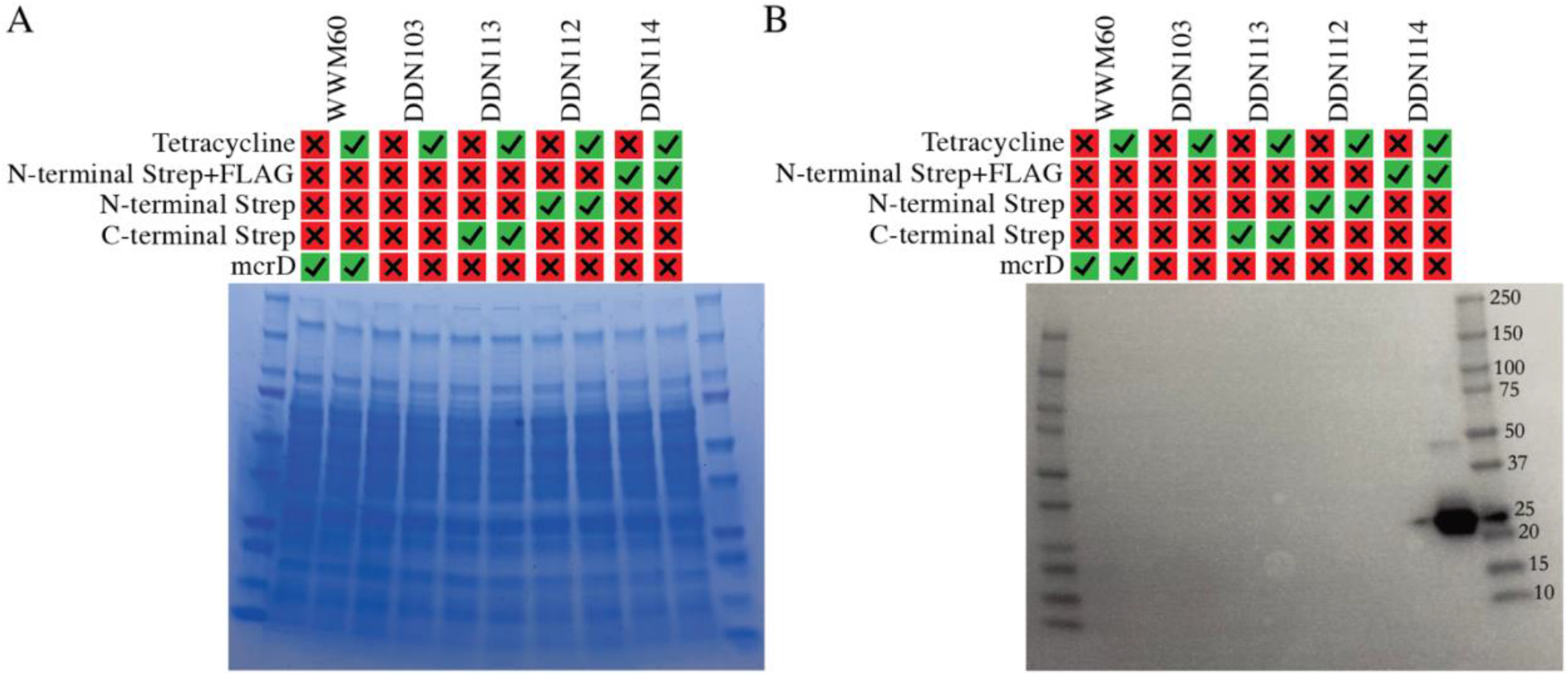
Expression of McrD *in trans*. **(A)** Coomassie and **(B)** anti-FLAG Western blot of crude protein extracts from *M. acetivorans* strains. Only strain DDN114, which contains a plasmid encoded *mcrD* with a N-terminal twin FLAG and Strep tags showed immunolabelling at the expected size. The plasmid encoded *mcrD* is under the control of a tetracycline-inducible promoter and 100 μg/mL tetracycline was added to the growth medium to induce expression of the tagged McrD protein.

**fig. S6.**
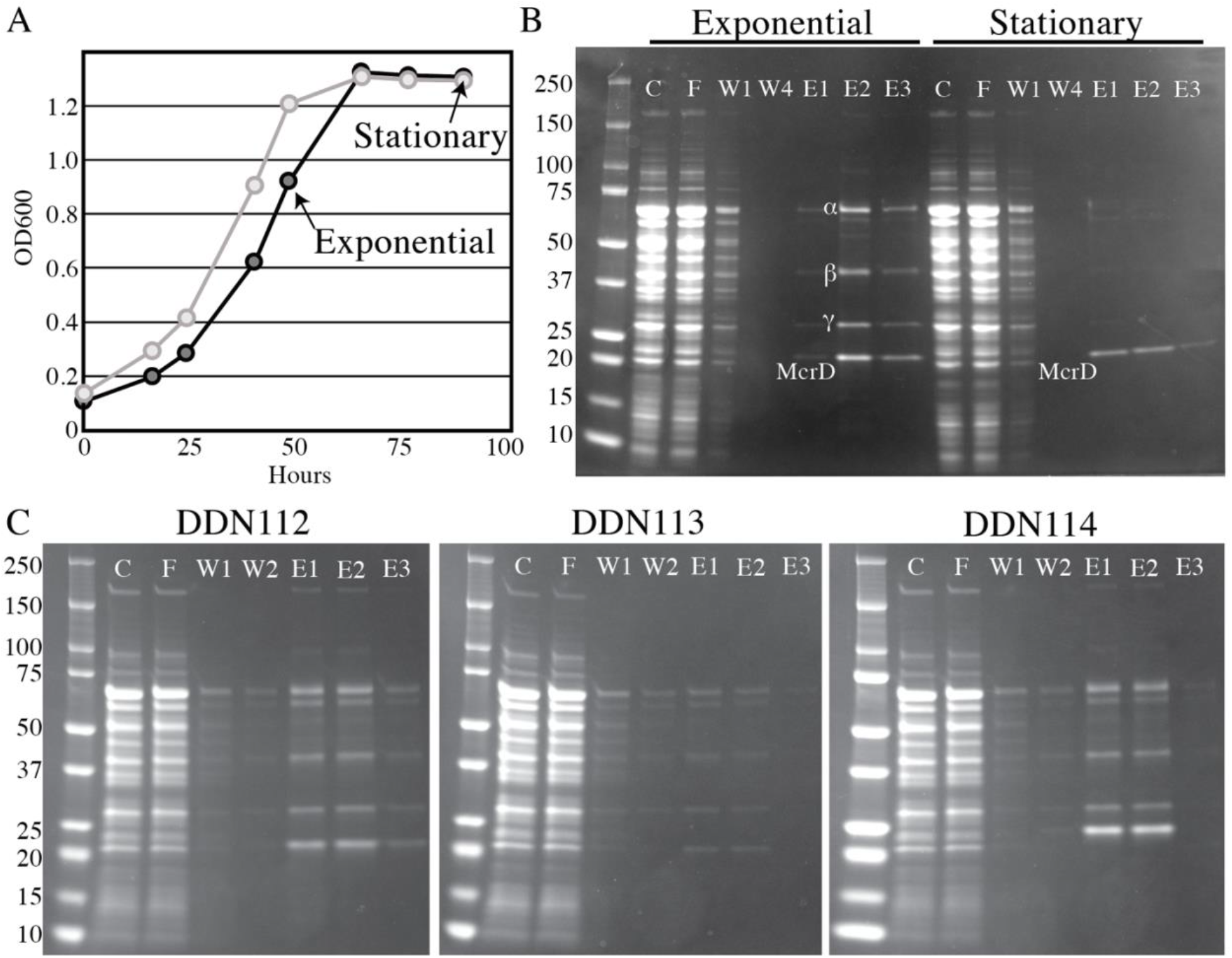
Purification of the McrABGD complex. (**A**) 500 mL cultures of DDN114 were grown and harvested in late exponential (dark gray line) or stationary phase (light gray line) for purification for the McrABGD complex. Optical density readings displayed here were taken from Balch tubes set up in parallel to the large format bottle incubations to determine the growth phase. Arrows indicate the timepoint where the 500mL cultures were harvested. (**B**) SDS-PAGE gels showing Crude cell lysate (C), flow-through of Strep-Tactin Superflow Plus resin used for affinity purification (F), first wash (W1), fourth and final wash (W4), first, second and third elution (E1, E2 and E3, respectively). Protein purification from exponential phase demonstrate roughly stoichiometric amounts of subunits α, β, γ and δ, while cells in stationary phase yield almost entirely δ free from other proteins. **C**) Comparison of tags on the co-purification of MCR subunits. DDN112 (N-terminal Strep only), DDN113 (C-terminal Strep only) and DDN114 (N-terminal FLAG and Strep) all pulled down α, β and γ subunits along with the dominant δ band. Note: the band below the α subunit was identified as Hsp60 (MA0086) but was not detected in the cryoEM data (data not shown).

**fig. S7.**
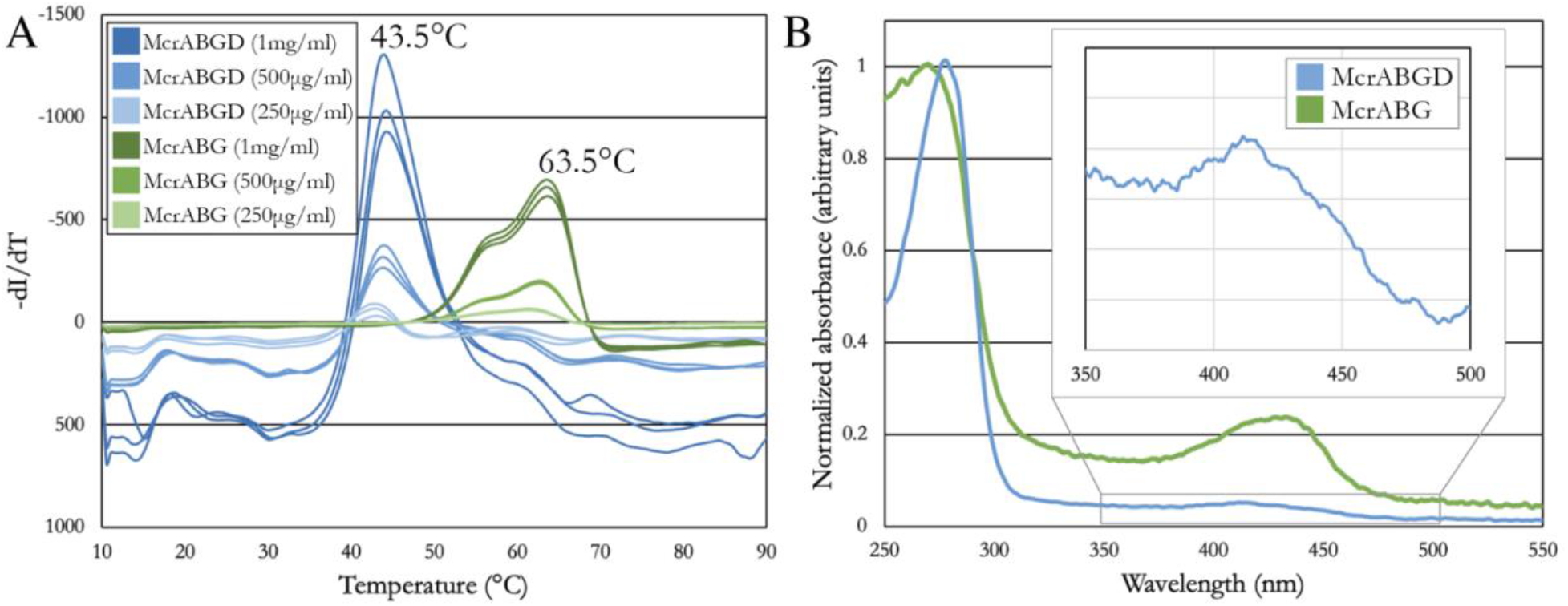
Properties of McrABGD complex. (**A**) SYPRO Orange melt curve analysis of McrABGD complex purified from DDN114 reveals a 20°C lower melting temperature than McrABG purified from WWM1086. (**B**) Absorbance spectra of McrABGD complex purified from DDN114 and McrABG purified from WWM1086. Inset region of showing weak absorbance between 400 and 450 nm consistent with sub-stoichiometric loading of McrABGD complex with cofactor F_430_.

**fig. S8.**
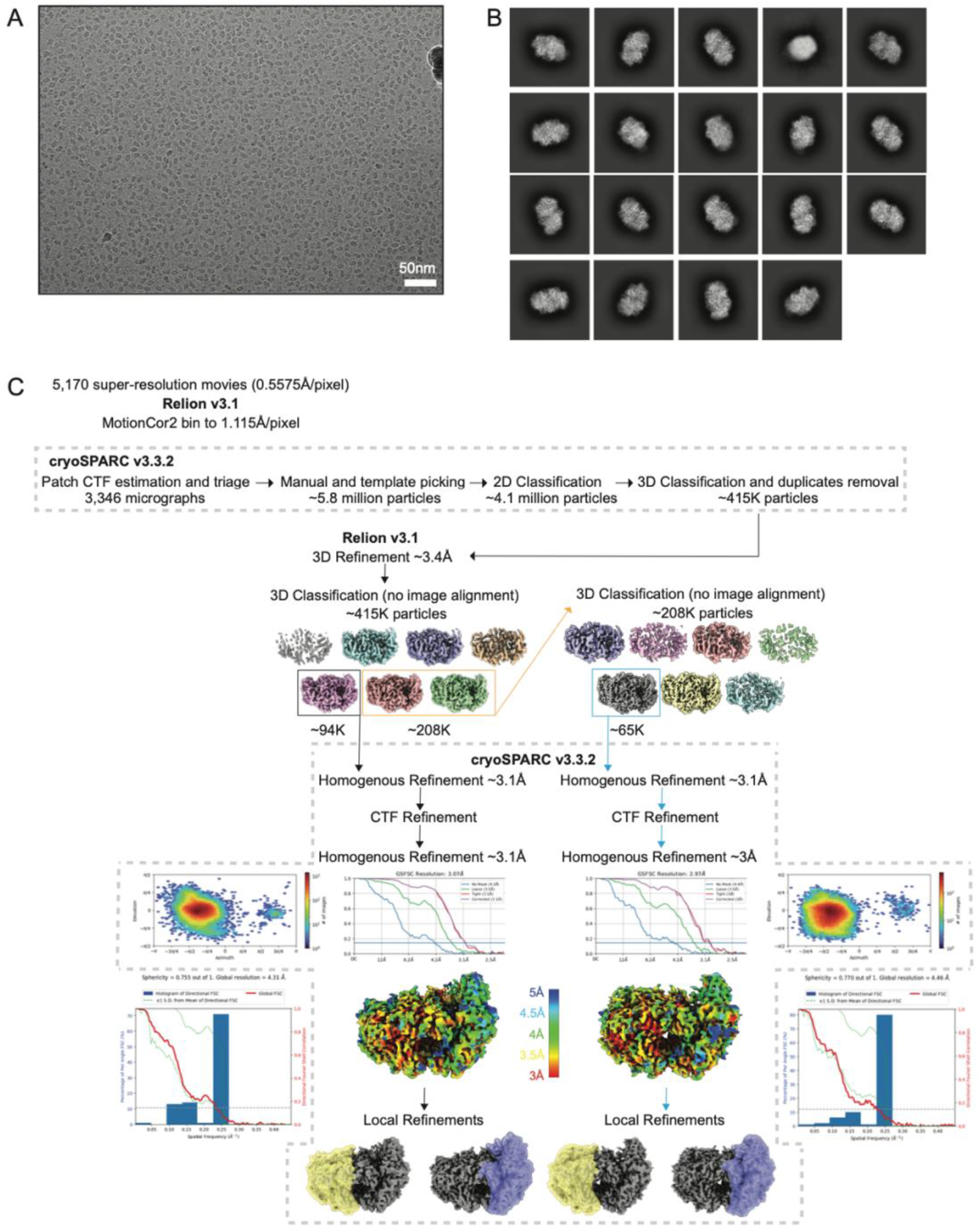
Cryo-EM processing pipeline. (**A**) Example micrograph of the McrABGD complex at - 2.1 μm estimated average defocus. Scale bar, 100Å. (**B**) Representative 2D class averages generated in cryoSPARC. The edge of each box corresponds to 250Å. (**C**) EM density from a well-resolved region of the final 3D reconstructions. (**D**) Flow chart depicting the steps taken during data processing to generate the two distinct conformational reconstructions of McrD-bound MCR complex. Steps enclosed in gray dashed boxes were performed in cryoSPARC. Sphericity was calculated using the 3DFSC script from D. Lyumkis, available at https://3dfsc.salk.edu.

**fig. S9.**
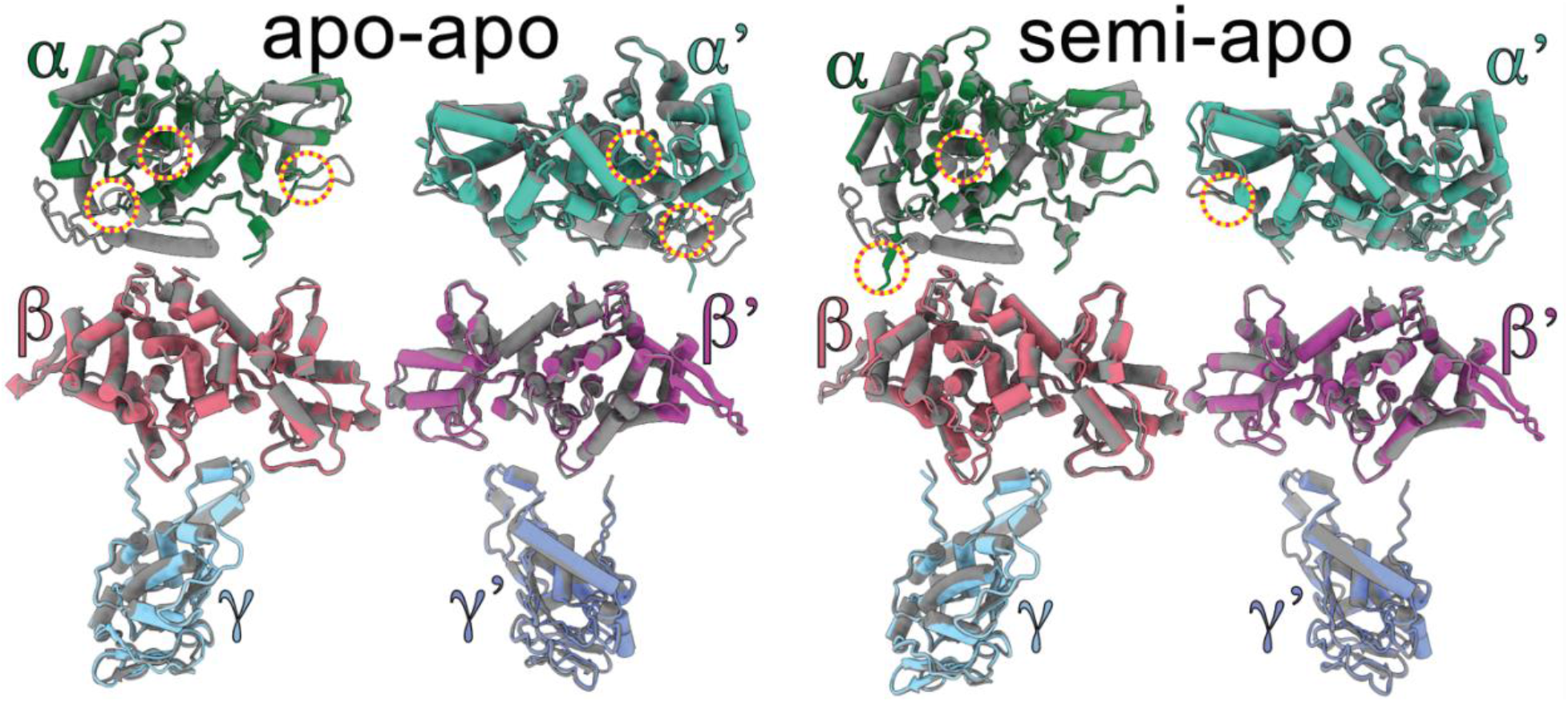
Details of individual chains. Each individual subunit of MCR from the cryoEM models reported here are aligned to their corresponding subunit from PDB:1e6y. Overlays highlight the extensive structural similarity, while major differences are highlighted in yellow dashed circles.

**fig. S10.**
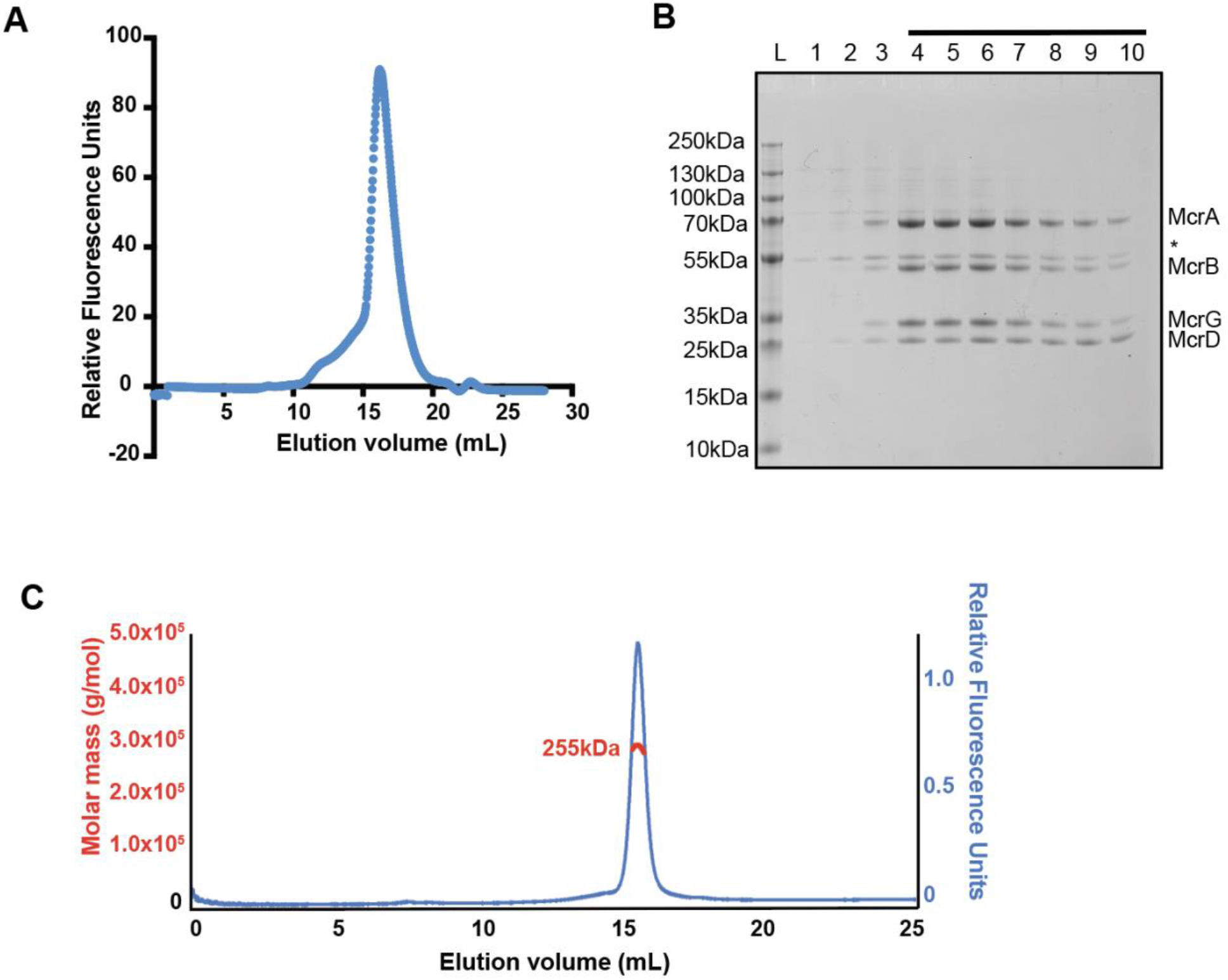
Size of the McrABGD complex and sample preparation for CryoEM. **(A)** Representative UV trace from gel filtration of the McrABGD complex on a Superose 6 Increase 10/300 GL. (**B**) Coomassie-stained gel corresponding to the peak fractions in **A**, * indicates multimeric McrD complex due to incomplete denaturating conditions in the gel (McrD verified by mass spectrometry, data not shown). (**C**) Representative trace of the purified McrABGD complex after gel filtration from SEC-MALS experiment.

**fig. S11.**
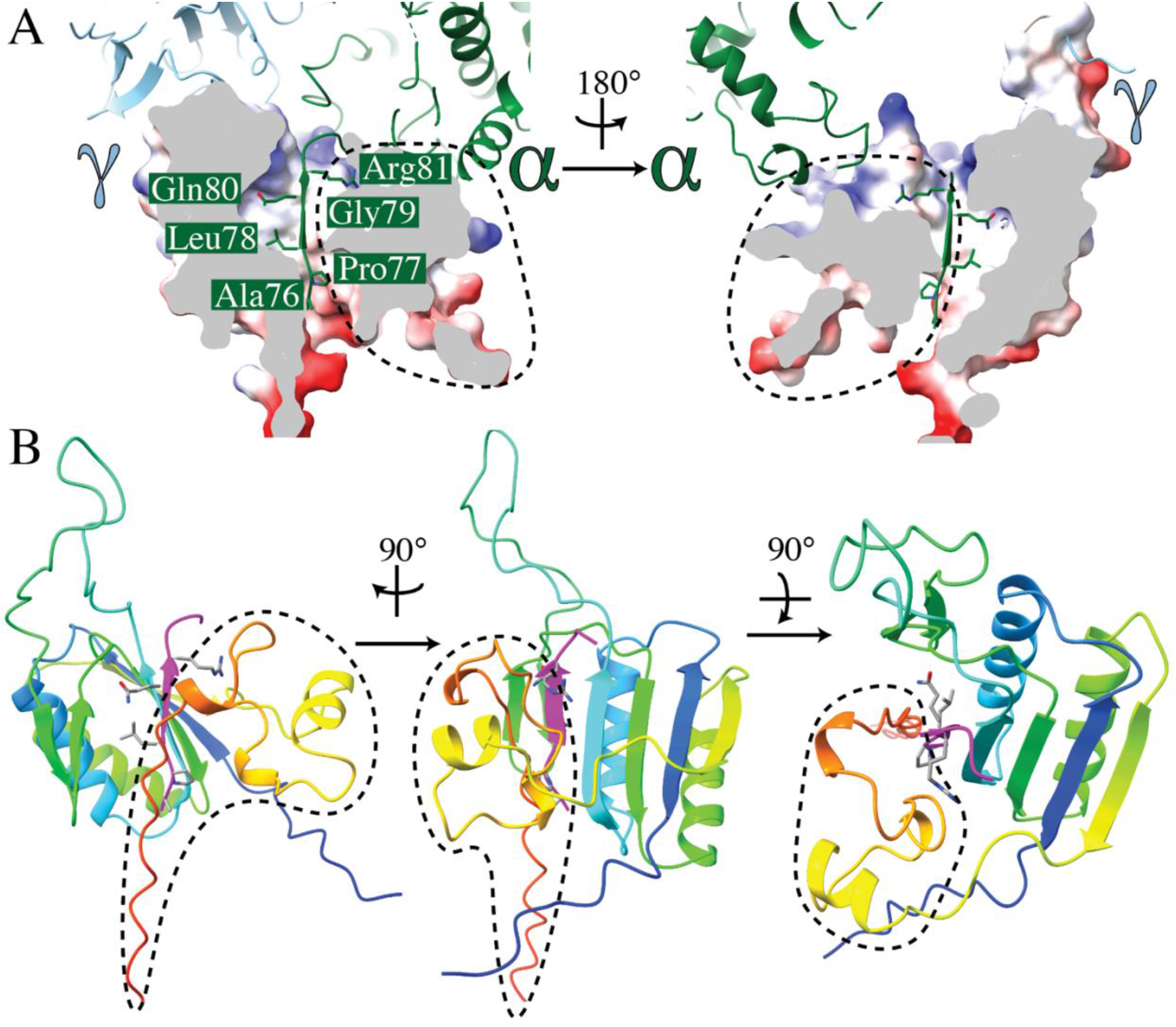
Unresolved C-terminal domain of McrD. The C-terminal domain of McrD past Asp^D129^ does not have corresponding CryoEM density (Tyr ^D130^-Glu^D170^). The AlphaFold2 prediction of tagged McrD which matches the N-terminal domain nearly perfectly predicts these C-terminal residues to fold in a way that produces a channel into which the residues Ala^α76^-Arg^α81^ fit neatly. (**A**) Two views of this interface with a cartoon diagram of the α and γ subunits and surface cutaway of the AlphaFold2 prediction of McrD. Side chains for Ala^α76^-Arg^α81^ are shown to highlight the fit. (**B**) Three views of the same interface with the McrD prediction shown as a cartoon with rainbow coloring. In all panels region of McrD not found in our cryoEM density is outlined with a dashed line.

**fig. S12.**
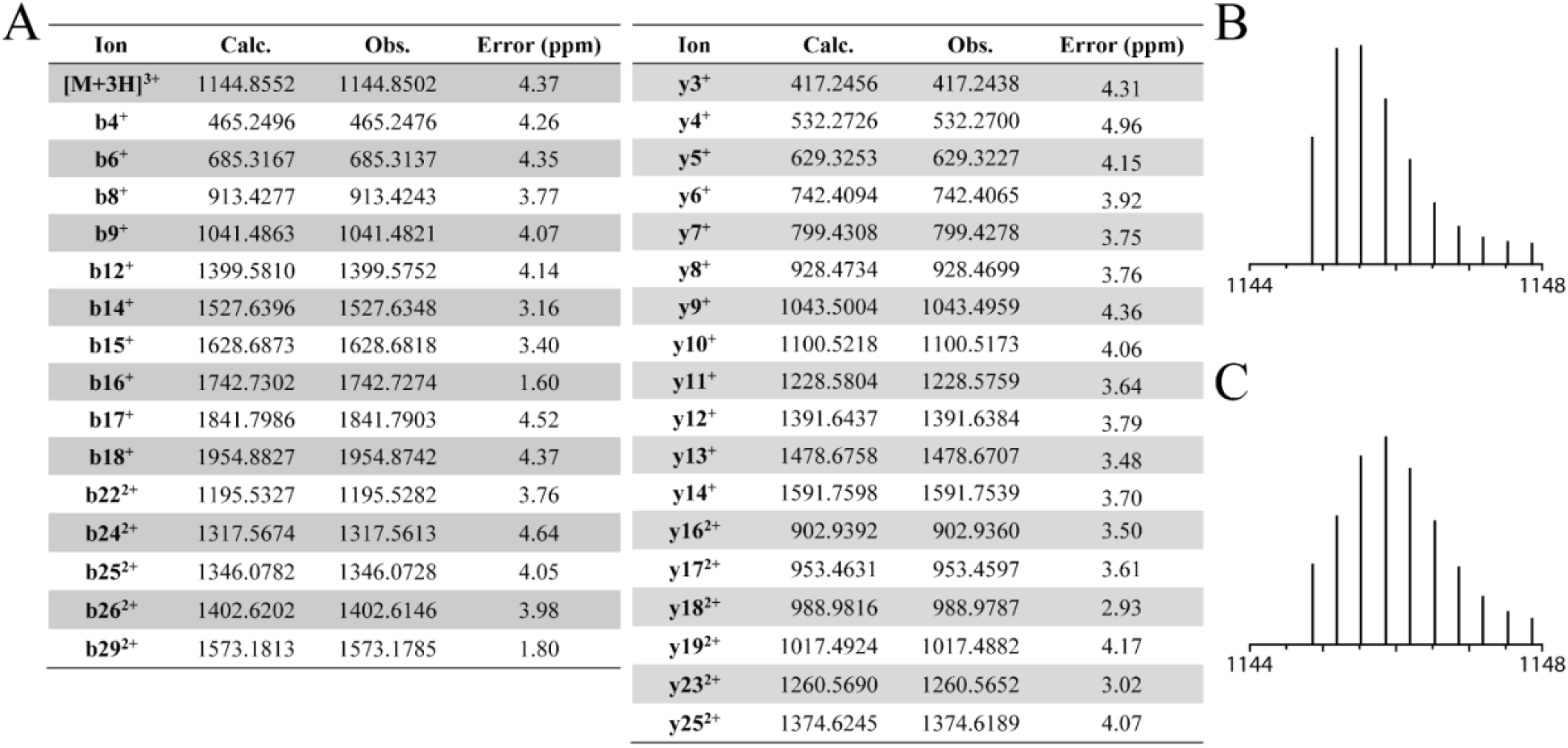
HRMS/MS analysis of an McrA tryptic peptide (Leu^461^-Arg^491^, m/z 1144.85). **(A)** Table of daughter ion assignments for an McrA tryptic peptide confirming three PTMs-thio Gly^465^, didehydro Asp^470^ and methyl Cys^472^ for fragmentation and annotated CID spectrum shown in Fig. 2D. Calculated isotope distribution for the parent ion **(B)** with thio Gly^465^, dehydro Asp^470^ and methyl Cys^472^ **(C)** 1:1 mixture of tryptic peptides having or lacking dehydro Asp^470^and containing thio Gly^465^ and methyl Cys^472^. The observed isotope distribution of the parent ion in **Fig. 2D** is similar to that in **(C)** suggesting that Asp^470^ is unmodified in some fragments.

**fig. S13.**
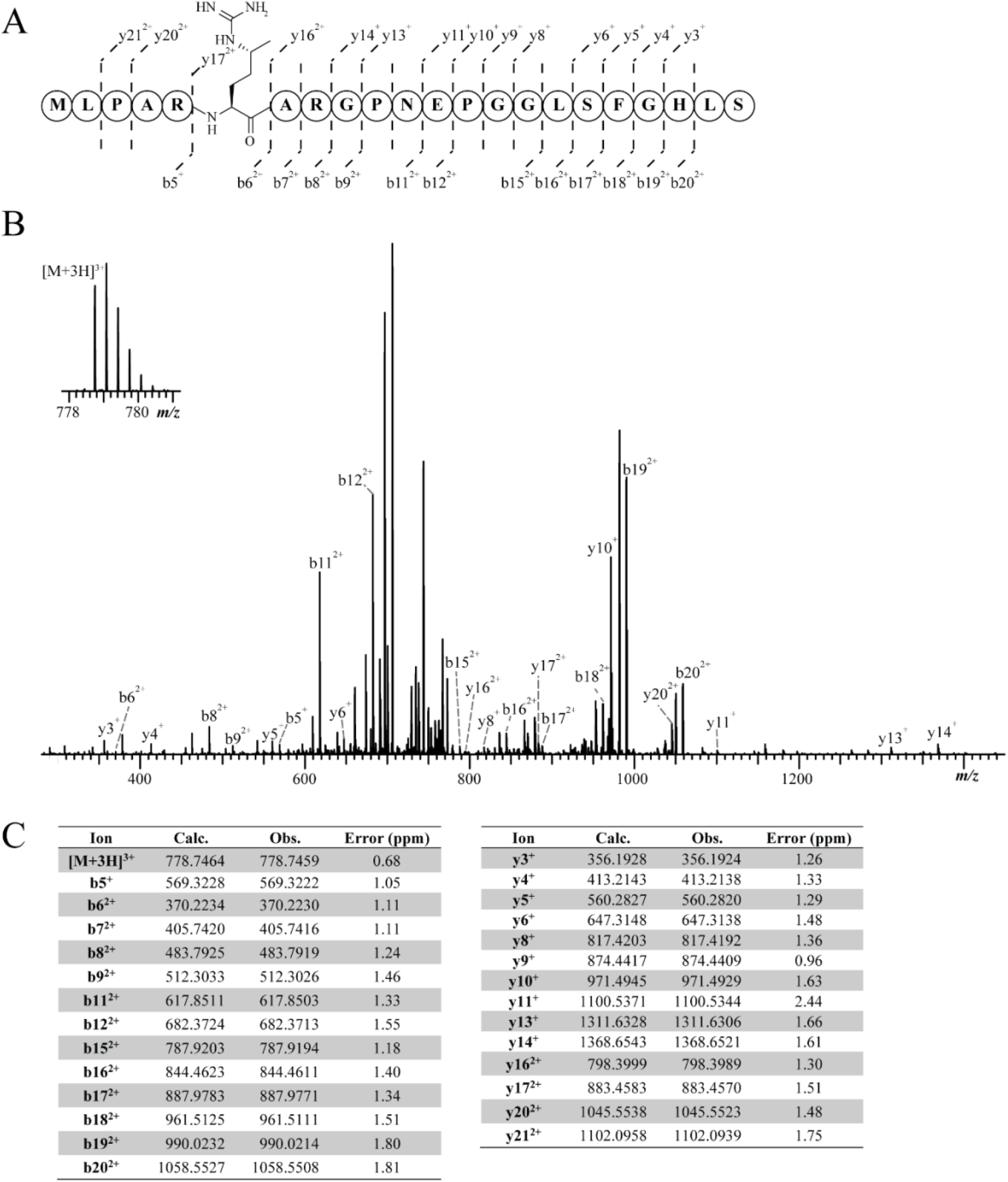
HRMS/MS analysis of an McrA AspN-GluC double-digest peptide (Met^280^-Ser^301^, m/z 778.74). **(A)**Fragmentation **(B)** Annotated CID spectrum of an McrA AspN-GluC double-digest peptide confirming methyl Arg^285^. Observed b and y ions are annotated. **(C)** Table of daughter ion assignments.

**fig. S14.**
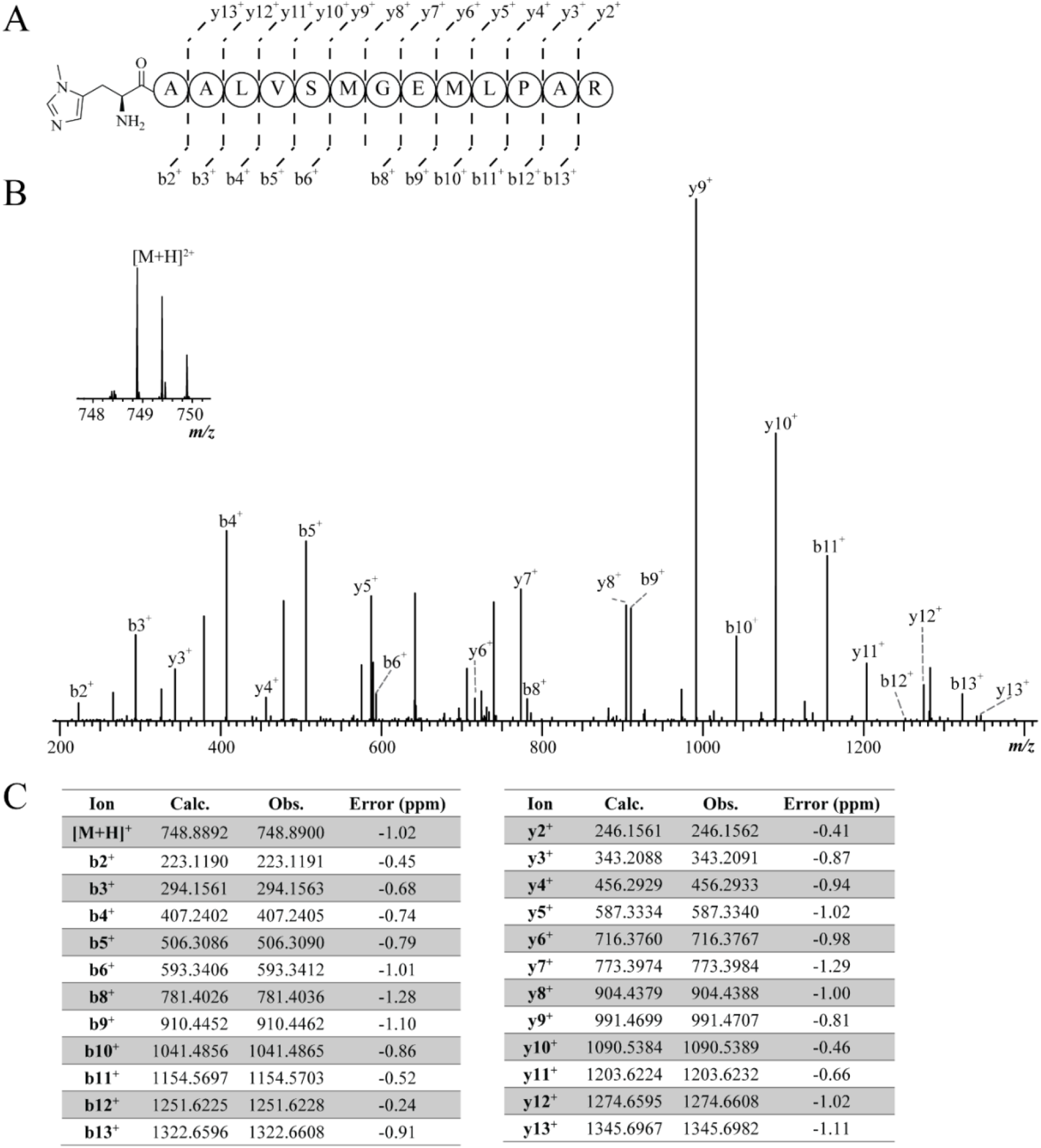
HRMS/MS analysis of an McrA tryptic peptide (His^271^-Arg^285^, m/z 748.88). **(A)** Fragmentation **(B)** Annotated CID spectrum of an McrA tryptic peptide confirming *N*-metlivl His^271^. Observed b and y ions are annotated. **(C)** Table of daughter ion assignments.

**fig. S15.**
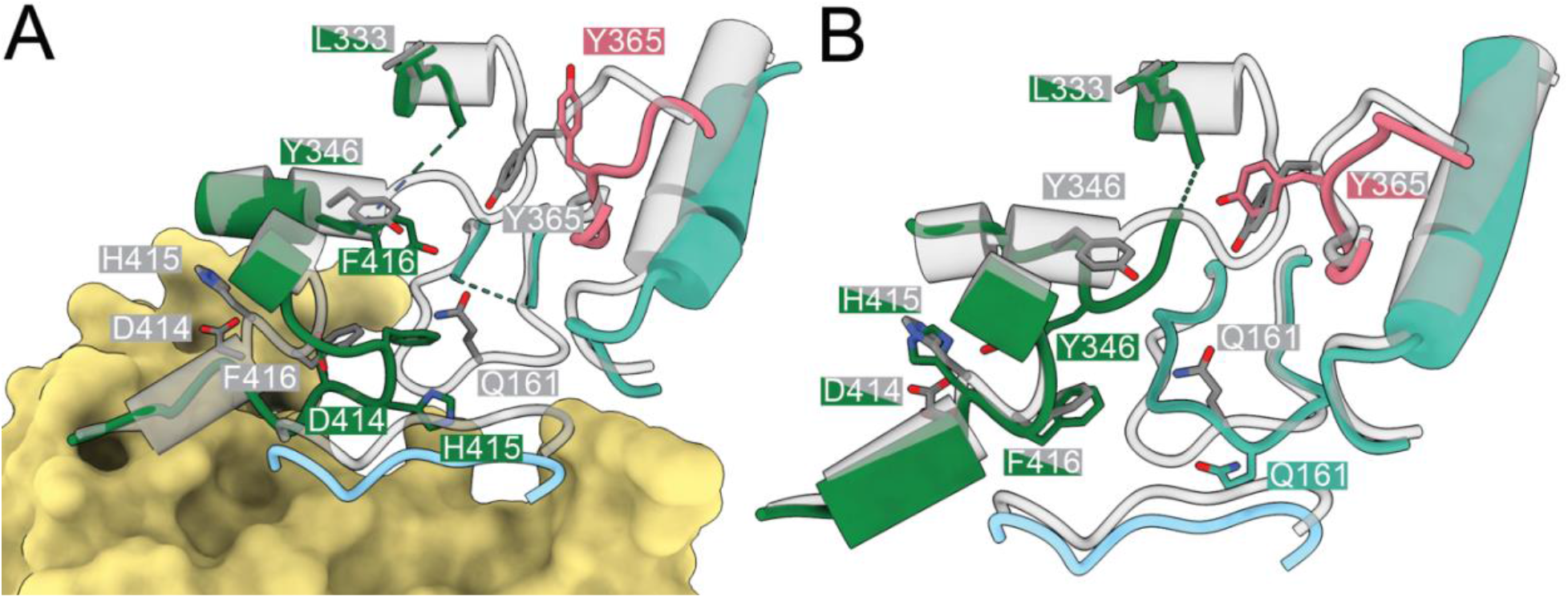
Active site details of apo-apo model. (**A**) McrD-bound active site in the apo-apo model shows similar deformations compared to that in the semi-apo model (**Fig. 3D**). (**B**) The contralateral active site in the apo-apo model shows a unique conformation, with notable displacements to the F_430_ axial ligand Q^α’161^ and a partially resolved disordered loop between Y^α346^ and L^α333^.

**Supplementary Table 1:**
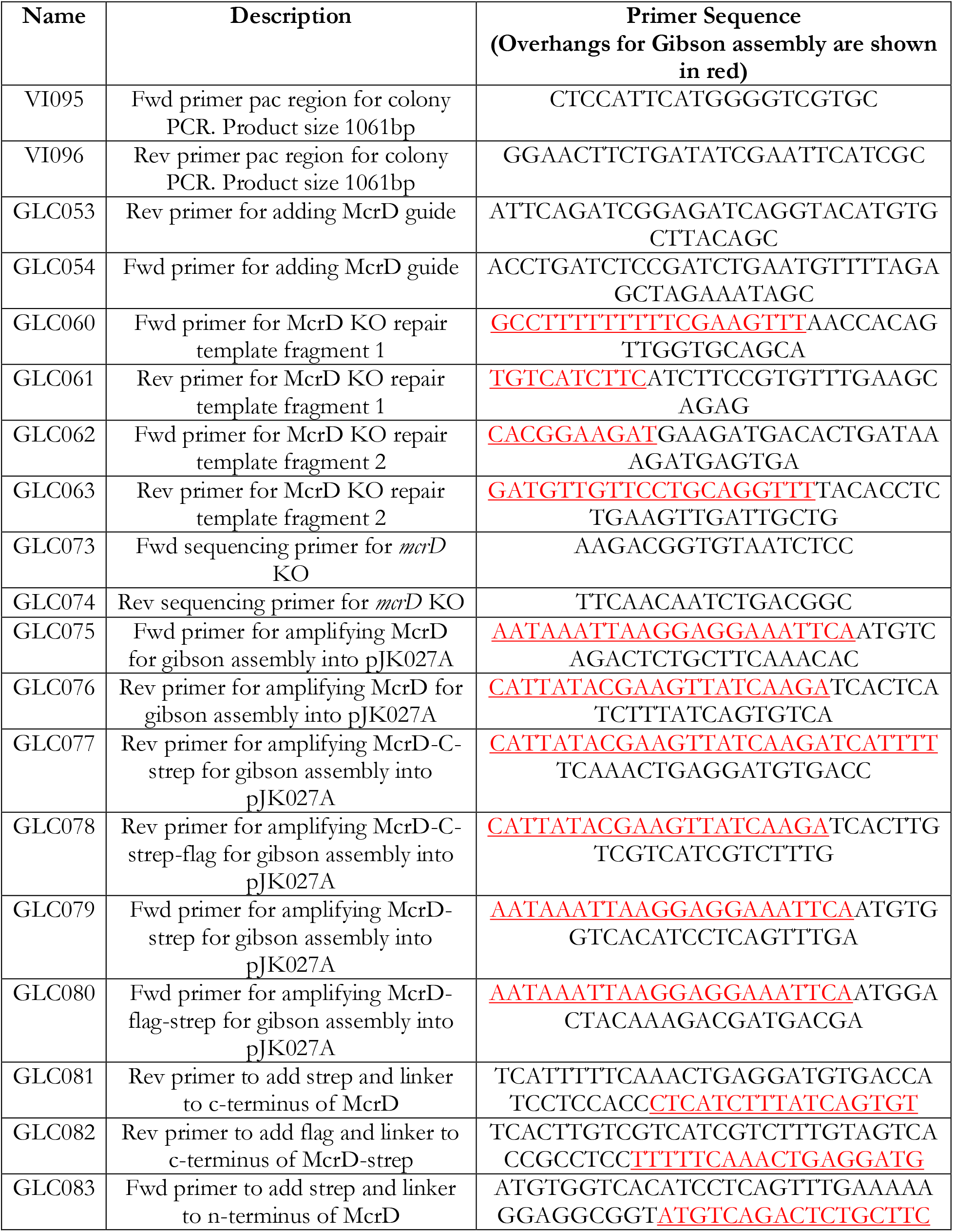
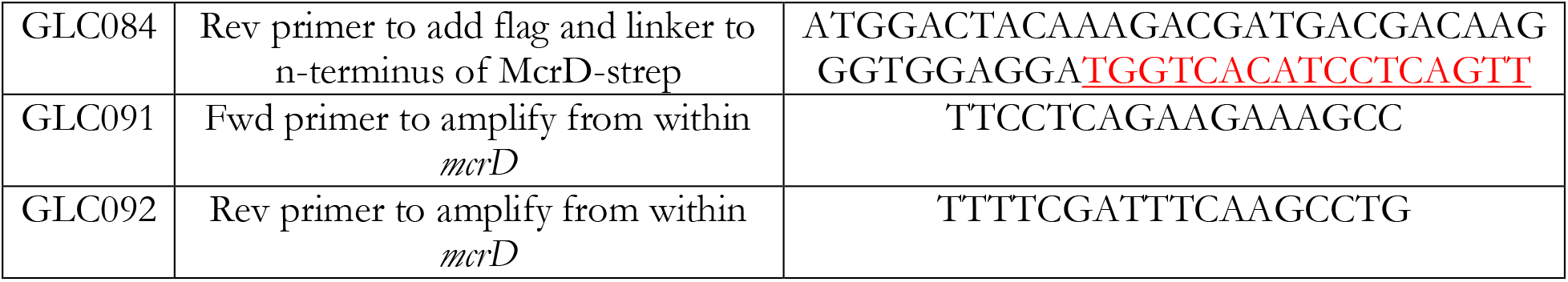
List of primers used in this study.

**Supplementary Table 2:** List of plasmids used in this study.

| Plasmid | Features | Source |
| --- | --- | --- |
| pAMG40 | Vector for fosmid retrofitting that contains pC2A and λattB | (Guss *et al.* 2008) |
| pJK027A | Vector with P*mcrB*(*tetO1*) promoter fusion to *uidA* that contains φC31-attB and λattP | (Guss *et al.* 2008) |
| pDN201 | pJK027A-derived plasmid with P*mcrB*(*tetO1*) promoter fusion to Spy *cas9* | (Nayak and Metcalf, 2017) |
| pGLC009 | pDN201-derived plasmid with synthetic fragments containing PmtaCB1 promoter fusion to sgRNA targeting the *mcrD* (*MA4549*) CDS | This study |
| pGLC011 | pGLC009-derived plasmid containing a repair template to generate an in-frame deletion of the *mcrD* (*MA4549*) CDS | This study |
| pGLC013 | Cointegrate of pGLC011 and pAMG40 | This study |
| pGLC023 | pJK027A-derived plasmid with P*mcrB*(*tetO1*) promoter fusion to *mcrD* (*MA4549*) CDS from *M. acetivorans* | This study |
| pGLC024 | pJK027A-derived plasmid with P*mcrB*(*tetO1*) promoter fusion to *mcrD* (*MA4549*) CDS from *M. acetivorans* with a C-terminal Strep tag | This study |
| pGLC026 | pJK027A-derived plasmid with P*mcrB*(*tetO1*) promoter fusion to *mcrD* (*MA4549*) CDS from *M. acetivorans* with a N-terminal Strep tag | This study |
| pGLC027 | pJK027A-derived plasmid with P*mcrB*(*tetO1*) promoter fusion to *mcrD* (*MA4549*) CDS from *M. acetivorans* with a N-terminal tandem Strep and FLAG tags | This study |
| pGLC028 | Cointegrate of pGLC023 and pAMG40 | This study |
| pGLC029 | Cointegrate of pGLC024 and pAMG40 | This study |
| pGLC031 | Cointegrate of pGLC026 and pAMG40 | This study |
| pGLC032 | Cointegrate of pGLC027 and pAMG40 | This study |

**Supplementary Table 3:** List of *Methanosarcina acetivorans* strains used in this study.

| Strain | Genotype | Source |
| --- | --- | --- |
| WWM60 | $\Delta hpt::P_{mcrB-tetR}$ | (Guss *et al.*, 2008) |
| WWM1086 | $\Delta hpt::P_{mcrB-tetR}$, Enterokinase cleavable TAP-tag at N-terminus of *mcrG* | (Nayak *et al.*, 2020) |
| DDN103 | $\Delta hpt::P_{mcrB-tetR}$, $\Delta mcrD$ | This study |
| DDN111 | DDN103/pGLC028<br>[*P_{mcrB(tetO1)-mcrD}*] | This study |
| DDN112 | DDN103/pGLC029<br>[*P_{mcrB(tetO1)-mcrD-C-terminal strep tag}*] | This study |
| DDN113 | DDN103/pGLC031<br>[*P_{mcrB(tetO1)- N-terminal strep tag-mcrD}*] | This study |
| DDN114 | DDN103/pGLC032<br>[*P_{mcrB(tetO1)- N-terminal strep and FLAG tag -mcrD}*] | This study |

**Supplementary Table 4:**
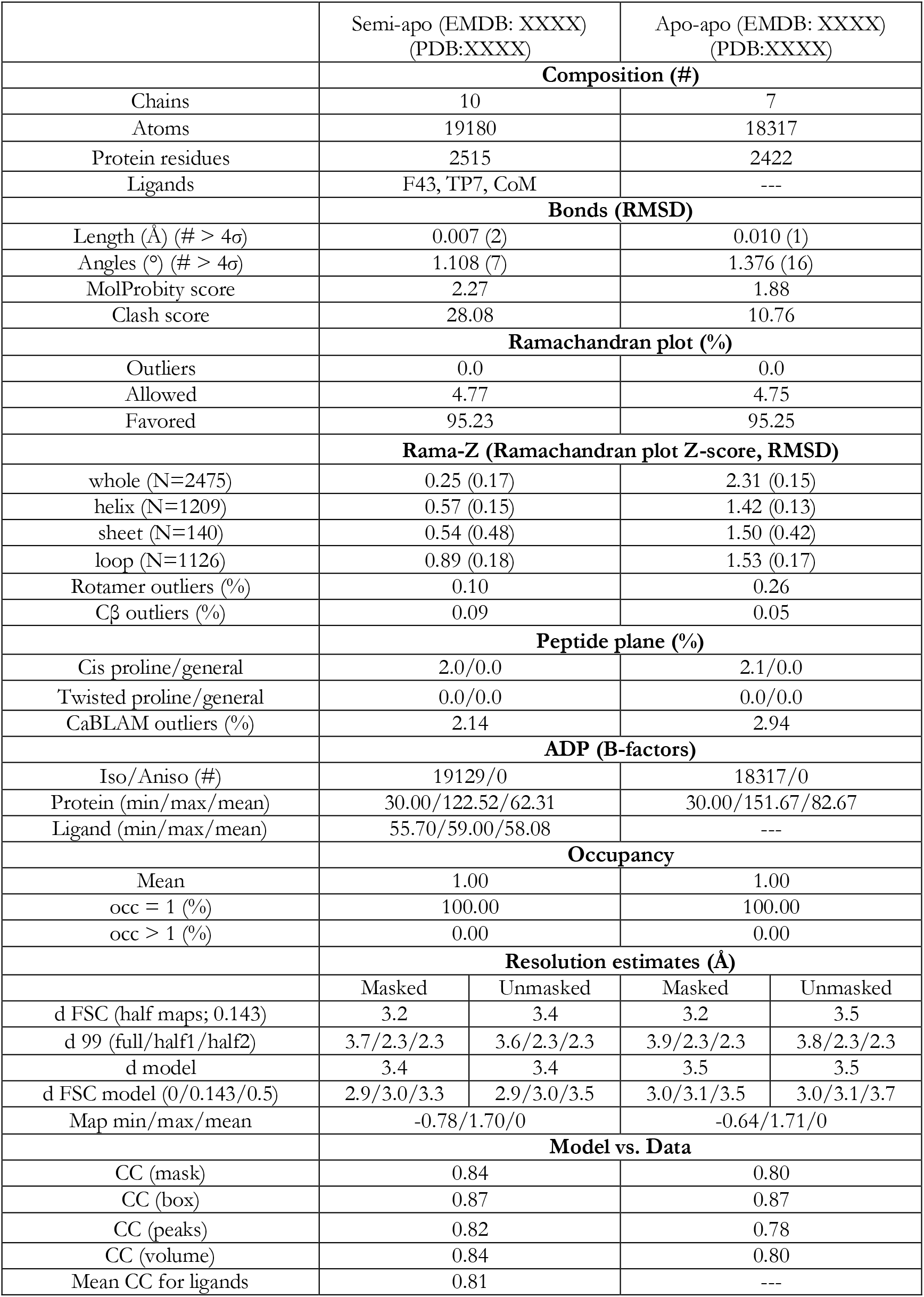
CryoEM structure statistics reported in this study.

## Supplementary Methods

### sgRNA design and plasmid construction

The Cas9 single guide RNA (sgRNA) sequence used in this study for genome editing was ACCTGATCTCCGATCTGAAT**<u>AGG</u>** (PAM sequence bold and underlined) and was generated with Geneious Prime v. 11.0. This guide targets the *mcrD* gene (MA4549) and matches 5,600,053-5,600,075 (+strand) in the *M. acetivorans* C2A genome. The CRISPR site finder tool was used with an NGG PAM site at the 3’ end with no off-target matches to the *M. acetivorans* genome allowed. The plasmids and primers used in this study are listed in Supplementary Tables 1 and 2 and were generated as previously described (*1*).

### Growth media and mutant generation

*M. acetivorans* strains were routinely maintained in 10ml of high salt (HS) media with either acetate (40mM), methanol (125mM) or trimethylamine (50mM) as the sole carbon and energy source as previously described (*2*). For transformation, 10ml cultures in late exponential phase were anaerobically pelleted and subjected to liposome-mediated transformation using establish protocols (*3*). Selection for transformants was carried out on HS media with 50 mM trimethylamine as the carbon source solidified with 1.5% agar and puromycin at 2μg/ml for positive selection. Colonies were screened for the desired mutation (see below) and restruck on plates with 20μg/ml 8ADP for counter selection to remove the genome editing plasmid and generate the clean deletion strain. Strains carrying tetracycline-inducible *mcrD* variants are permanently maintained in 2μg/ml puromycin to retain the expression plasmid. *M. acetivorans* strains used and generated in this study are listed in Supplementary Table 3.

### Mutant validation

PCR screening of colonies was carried out on whole cells resuspended in nanopure water amplified with GoTaq^®^ Green Master Mix (Promega Corporation). Primers were designed to amplify across the McrD gene (GLC073/74) and from within the McrD gene (GLC073/92 and GLC074/91). Positive identification of *ΔmcrD* genotype with no wild type contamination came from the expected decrease in product size with the 73/74 primer pair, and no product with the 73/92 or 74/91 primer pairs. PCR primers amplifying the Pac gene (VI095/96) were used to validate the loss of genome editing plasmids by 8ADP counter selection, or the presence of heterologous expression plasmids. Whole genome resequencing of the *ΔmcrD* strain was carried out by extracting genomic DNA from 10ml of stationary phase cuture grown in HS media with trimethylamine using the Qiagen blood and tissue kit (Qiagen, Hilden, Germany). Illumina library preparation and sequencing was carried out at SeqCenter (Pittsburgh, PA). Comparison of the *ΔmcrD* strain to *M. acetivorans* C2A was carried out using the breseq software package (*4*).Raw reads are deposited in the Sequencing Reads Archive (SRA) and are available under the BioProject XXXXXXXX.

### Growth measurements

*M. acetivorans* doubling times were determined by measuring the optical density at 600nm of cultures grown in Balch tubes containing 10ml HS media with growth substrates and media additions as indicated. Measurements were made using a UV-Vis spectrophotometer (Gensys 50, Thermo Fisher Scientific, Waltham, MA). Doubling times were determined by fitting all points in for which the best fit line of the log2 transformed optical density data had >0.99 R^2^ values. For Ni limitation experiments, trace metal solutions were prepared without Ni addition. For oxygen exposure experiments, the indicated volume of lab air was added through a 0.2 μm syringe filter (CellTreat Scientific Products, Pepperall, MA).

### RNA extraction and bioinformatic analysis of transcriptome

WWM60 and DDN103 were grown in triplicate in methanol media at 37°C until late log phase (~0.6 OD600). Two ml of culture was sampled and added immediately to equal amounts 37°C Trizol solution (Life Technologies, Carlsbad, CA, USA). After a 5 minute incubation at room temperature 4 ml of ethanol was added to the culture and Trizol mixture. Solution was applied to a Qiagen RNeasy Mini Kit (Qiagen, Hilden, Germany) and extraction proceeded according to the manufacturer’s instructions. DNAse treatment, rRNA depletion and Illumina library preparation and sequencing was carried out at SeqCenter (Pittsburgh, PA). Analysis of transcriptome data was carried out on the KBase bioinformatics platform (*5*). Briefly, raw reads were mapped to the *M. acetivorans* C2A genome using Bowtie2 (*6*), assembled using Cufflinks (*7*), and fold changes and significances values were calculated with DESeq2(*S*). Raw reads are deposited in the Sequencing Reads Archive (SRA) and are available under the BioProject XXXXXXXX.

### Protein extraction and affinity purification

For protein purification 500ml cultures of *M. acetivorans* strains were grown in HS media with trimethylamine (100mM). Cells were lysed and cell debris removed as previously described (*9*), with the one modification that the 50mM sodium phosphate was replaced with 50mM Tris HCl (pH 7.0) in all buffers. Strep-tagged proteins were purified using Strep-Tactin Superflow plus resin (Qiagen, Hilden, Germany) as described previously (*9*) again with the sole modification that sodium phosphate buffer was replaced with Tris HCl. Strep-tagged protein complexes were eluted with washing buffer containing 2.5 mM desthiobiotin.

### Protein gels

Protein preparations were mixed with 4x Laemmli sample buffer (Bio-Rad, Hercules, CA) and final concentration of 355mM 2-mercaptoethanol and heated at 90°C for 5 minutes before being run on Mini-Protean TGX denaturing SDS-PAGE gel (Bio-Rad, Hercules, CA). Coommassie staining was carried out with GelCode™ Blue Safe Protein Stain (Thermo Fisher Scientific). For immunoblotting, gels were transblotted using the Trans-Blot Turbo transfer system (Bio-Rad, Hercules, CA) onto 0.2 μm PVDF membrane (Bio-Rad, Hercules, CA). Monoclonal anti-Flag M2-Peroxidase (HRP) antibody (Sigma-Aldrich, St Louis, MO) was used at 50,000 fold dilution as previously described (*1*). Immobilon Western Chemiluminescent HRP Substrate (Millipore, Burlington, MA) was used with manufacturer’s protocol for detecting HRP-conjugated antibodies. Coomassie and immublots were imaged with the ChemiDoc MP Imaging System (Bio-Rad, Hercules, CA).

### Proteolytic digestion of purified McrABDG complex

For in-solution trypsinolysis (His^α_2_71^-Arg^α_2_85^, m/z 748.88), 50 μg of purified McrABDG complex was digested with Sequencing Grade Modified Trypsin (1:100 w/w ratio; Promega) at a final McrABDG concentration of 0.5 mg/mL in 50 mM Tris-HCl, 1 mM CaCl2 at 37°C for 14 h. For in-gel trypsinolysis of McrA (Leu^α461^-Arg^α491^, m/z 1144.85), 100 μg of purified McrABDG complex was separated by SDS-PAGE. In-gel trypsin digest was performed per established protocols on excised gel pieces corresponding to McrA (*10*).

For AspN and GluC double digestion (Met^α_2_80^-Ser^α301^, m/z 778.74), 300 μg of purified McrABDG was digested with endoproteinase GluC (1:200 w/w ratio; Promega) and endoproteinase AspN (1:125 w/w ratio; Promega) at a final McrABDG concentration of 2 mg/mL in 50 mM NH_4_HCO_3_ pH 8 at 37°C for 14 h. MeCN was added to the resultant digest to a final ratio of 50/50 50 mM aq. NH_4_HCO_3_/MeCN and centrifuged at 17,000g to remove precipitate. The supernatant was dried using a SpeedVac concentrator prior to purification by HPLC using a Thermo Fisher Vanquish UHPLC equipped with a Thermo Scientific Accucore C18 column (150 × 4.6 mm, 2.6 μm particle size, 80 Å pore size). The dried sample was resuspended in 5/95 10 mM aq. NH_4_HCO_3_/MeCN and injected with a mobile phase of 10 mM aq. NH_4_HCO_3_/MeCN and separated using a gradient from 5-65% MeCN over 25 min at 1.5 mL/min. Fractions collected every min were analyzed by MALDI MS to identify those containing peptide fragment of interest. Desired fractions were dried using a SpeedVac concentration for subsequent HRMS/MS analysis.

### HRMS/MS analysis of McrA peptide fragments

HPLC-purified, dried samples were resuspended in ESI mix [80% MeCN, 19% H2O, 1% acetic acid]. Digests were desalted using ZipTips, eluted into 75% aq. MeCN, and diluted 1:1 in ESI mix before HRMS/MS analysis. Samples were directly infused onto a ThermoFisher Scientific Orbitrap Fusion ESI-MS using an Advion TriVersa Nanomate 100. MS calibration was performed with Pierce LTQ Velos ESI Positive Ion Calibration Solution (ThermoFisher). The MS was operated using the following parameters: 120,000 resolution, 0.4-2 m/z isolation width (MS/MS), 35 normalized collision energy (MS/MS), 0.4 activation q value (MS/MS), and 30 ms activation time (MS/MS). Fragmentation was performed using collision-induced dissociation (CID) at 30%, 50%, and 70%. Data analysis was conducted using the Qualbrowser application of Xcalibur software (ThermoFisher Scientific).

### SEC MALS

A Superose6 10/300 Increase column (Cytiva) was attached in line to an Agilent Technologies 1100 series with a 1260 Infinity lamp, Dawn Heleos II and the Optilab T-Rex (Wyatt Technology) operating on Astra v5.3 software (Wyatt Technology). The system was equilibrated overnight in 25mM Tris pH 8, 150 mM NaCl before standardization with 2mg/mL bovine serum albumin. 125uL of McrABGD sample was injected onto the system at 0.35mL/min and light scattering was used to determine the molecular weight.

### Sample preparation for cryoEM

Frozen aliquots of protein were thawed and centrifuged at 21,000g for 5 minutes before gel filtration chromatography on an Akta Purifier (GE) equipped with a Superose6 10/300 Increase column (Cytiva) into GF buffer (25mM Tris pH 8, 150mM NaCl). The peak fractions (see Supplemental Figure) were collected and concentrated in a 100K MWCO spin concentrator (Millipore). 3uL of sample at 0.55mg/mL was deposited onto freshly glow-discharged Quantifoil R 2/1 Copper 300 mesh grids (Electron Microscopy Sciences, CAT# Q310CR1) for 5 seconds before being blotted at 4°C under 100% humidity for 3 seconds in a Vitrobot Mark IV (ThermoFisher) at blot force 18 and plunged into liquid ethane. The grids were glow discharged beforehand for 30s at 25mA at 0.37mBar in a PelCo EasiGlo (Ted Pella, Inc.)

### CryoEM data collection

~5,200 super-resolution dose-fractionated images were collected with a total dose of 50^e-^ over a defocus range of −0.8um to −2.0um on a Talos Arctica, equipped with a Gatan K3 camera, operating at 200kV with a nominal magnification of 36,000X and a pixel size of 0.5575Å/pixel. Using SerialEM (*11*), movies were collected in a 9-hole image shift pattern with 2 exposures captured per hole, for a total of 18 images per stage shift.

### CryoEM data processing

Movies were motion-corrected with 2x binning using MotionCor2 (*12*) via the RELION wrapper (*13*) and then imported into cryoSPARC (*14*) for patch CTF estimation, micrograph curation, and particle picking, which generated 5.7 million particles. Junk particles were filtered out via 2D Classification, yielding 4.1 million particles. Duplicate particles were removed, and the remaining particles were pruned via 3D classification, by retaining only the classes with obvious density for McrD, leaving 415K particles. This particle set was transferred into RELION via csparc2star.py (*15*), for 3D refinement to generate a consensus reconstruction followed by 3D classification without alignments. Two distinct classes emerged, one with 94K particles and another with 208K particles. The 208K particle stack went through a second round of 3D classification and ultimately yielded a stack with 65K particles. The separate 94K and 65K particle stacks were imported back into cryoSPARC for additional homogenous refinements, CTF refinement, and estimation of local resolution. 3DFSC (*16*) was used to measure the directional resolution anisotropy. Due to high anisotropy and variability in resolution, local refinements were performed using masks at each end of the molecule. The local refinement maps were combined into composite maps using Phenix (*17*). A mix of composite, sharpened, and local refinement maps was used for interpretation.

### Model building and refinement

A crystal structure of McrABG complex (pdb: 1e6y) was docked into the semi-apo map but left a sizeable patch of density unoccupied (*18*). An AlphaFold2 prediction of McrD was simultaneously placed into the remaining density (*19*), and together these models were used as the starting point for manual rebuilding throughout the cryoEM density using Coot (*20*). The resulting semi-apo model was docked into the apo-apo map and used as the initial model for manual rebuilding. The rebuilt models were iteratively refined in real space using Phenix, and figures were produced using ChimeraX (*21*).

## Notes

### Competing Interest Statement

The authors have declared no competing interest.

